# Diverse epigenomic mechanisms underpin transcriptional dysregulation in Polycomb-altered acute myeloid leukemia

**DOI:** 10.1101/2025.08.28.672871

**Authors:** Cosmin Tudose, Luke Jones, Theodora-Ioana Grosu, Marie-Claire Fitzgerald, Noura Maziak, Rebecca Ling, Anindita Roy, Juan M. Vaquerizas, Colm J. Ryan, Jonathan Bond

## Abstract

**Background:** Polycomb Repressive Complex 2 (PRC2) modulates chromatin accessibility and architecture to direct tissue-specific gene expression. PRC2 function is frequently altered in cancer by loss-of-function mutation or deletion, but the downstream effects on transcriptional regulation are incompletely understood.

**Results:** To gain insights into these mechanisms, we performed a holistic analysis of epigenomic and transcriptional changes in an isogenic model of acute myeloid leukemia (AML) with heterozygous *EZH2* deletion that mimics reduced PRC2 function in patient leukemias.

PRC2-depleted cells had diverse gene expression changes, including a bias towards more immature monocyte-lineage transcriptional signatures. PRC2 depletion also correlated with marked increases in chromatin accessibility genome-wide, with 10-45% increases in ATAC-seq peaks in EZH2+/− clones. These changes were accompanied by decreased H3K27me3 and increased H3K27ac levels in CUT+RUN assays that were incompletely linked to transcriptional activity.

Despite these generalised changes, 3D chromatin architecture assessed by Hi-C was largely maintained, with H3K27me3 preferentially lost in regions with low DNA-DNA contact frequency. Surprisingly, some regions gained broad H3K27me3 domains at heavily compacted chromatin. We notably saw compartmentalisation changes upstream of the transcriptionally upregulated fetal hematopoiesis gene *LIN28B* in EZH2+/− cells, with corresponding activation of a LIN28B-specific transcriptional program, including upregulation of the *CDK6* oncogene. These results correlated with EZH2+/− cell phenotype, including decreased cellular proliferation and increased resistance to CDK6 inhibitor palbociclib.

**Conclusions:** Our findings suggest that PRC2 depletion pleiotropically affects AML transcriptional regulation to directly impact cell phenotype and treatment responsiveness, which may partially explain the aggressive biology seen in these cases.

## Introduction

Polycomb group proteins (PcGs) act as multiprotein complexes to regulate lineage-specific gene expression by diverse mechanisms during normal development and cell fate commitment [1]. The primary functions of Polycomb Repressive Complexes (PRCs) are thought to be linked to chemical modification of histones at transcriptionally inactive loci, with PRC2 and PRC1 marking chromatin with histone 3 lysine 27 trimethylation (H3K27me3) and histone 2A lysine 119 ubiquitination (H2AK119ub), respectively [2,3]. Aside from their enzymatic activities, PRC2 and PRC1 shape the chromatin landscape by mediating long-range looping and by creating largely repressed polycomb-associated domains, also known as polycomb bodies [4–6]. In line with this long-range function, disruption of H3K27me3-mediated loop anchors can activate genes at megabase distance [7]. PRC1 is also important for correct nucleosome positioning at transcriptional start sites (TSSs) [8]. In contrast, the role of PRC2 in nucleosome positioning is less clear. While some evidence indicates that it has no role in the nucleosome landscape [8], EZH2 depletion has been shown to lead to altered nucleosome occupancy and increased chromatin accessibility at bivalent promoters (i.e., marked with H3K4me3 and H3K27me3) at certain loci [9].

Consistent with PcG roles as global transcriptional regulators, altered Polycomb function is associated with a range of constitutional and acquired pathologies [10]. In cancers, PcG changes are usually linked to loss of PRC function caused by somatic mutations and deletions, which frequently correlate with tumour biology and patient outcomes. For example, loss-of-function mutations and deletions in core PRC2 factors *EZH2*, *SUZ12*, and *EED* are commonly found in acute leukemias, predicting treatment responses in both acute myeloid leukemia (AML) and T-acute lymphoblastic leukemia (T-ALL) [11–13]. In the case of AML, *EZH2* mutations and low EZH2 expression have been shown to correlate with chemoresistance [12,14–16], but a mechanistic understanding of how altered PRC2 function affects leukemia phenotype is lacking.

It is logical that some of these effects are due to dysregulation of functions that PRC2 normally performs during blood cell development. For example, EZH2 is essential for maintaining hematopoietic stem and progenitor cell (HSPC) identity by mediating H3K27me3 placement and 3D chromatin architecture [17]. Furthermore, EZH2 is crucial in repressing fetal hematopoiesis programs in adult blood cells by repressing a specific transcriptional program controlled by the Let-7 miRNA suppressor LIN28B [18]. Understanding these processes in leukemia cells is however further complicated by some additionally reported effects of PcG proteins in this context. Despite its tumour suppressor function, we also know that PRC2 activity can be important in AML tumorigenicity and cross-talks with other frequently altered epigenetic components [19,20]. In a *KMT2A*-rearranged (*KMT2A*r) context, [21] and [22] showed that EZH2 can also act as an oncogene, with cells that have *KMT2A*::*MLLT3* translocations being dependent on EZH2 activity. However, in *KMT2A*r murine models, [15] showed that EZH2 acts as a tumour suppressor at disease induction, decreasing survival of mice with EZH2-depleted leukemia. Conversely, *EZH2* depletion on the same genetic background during AML maintenance results in better mouse survival [15].

To investigate how altered PRC2 function mechanistically affects transcriptional regulation in leukemia, we decided to create an isogenic cell line model of heterozygous *EZH2* deletion on a common AML genetic background. Integrated epigenomic and transcriptomic characterisation of this system reveals pleiotropic and diverse roles for PRC2 in regulating chromatin accessibility, transcription, and genome architecture in this context. Importantly, these changes could be linked directly to AML cell phenotype, providing clues as to how PRC2 alterations might ultimately influence leukemia biology.

## Results

### EZH2 depletion alters lineage-specific transcriptional programs

To model PRC2 loss-of-function in patient leukemia cells, we used CRISPR-Cas9 to edit *EZH2* in the myelomonocytic AML cell line OCI-AML2. To ensure that our experimental results would not be clone-specific [23], we generated two models of heterozygous EZH2 depletion, henceforth named clone 5 (C5) and clone 9 (C9) (Figure 1A, Supplementary Figure S1A). We confirmed decreases in EZH2 protein and H3K27me3 by approximately 20-40%, in line with what is seen in *EZH2*-mutated patient leukemia cells *in vivo* [14]. As expected, [24,25] H3K27ac levels were higher in EZH2-low cells. (Figure 1B).

**Figure 1.**
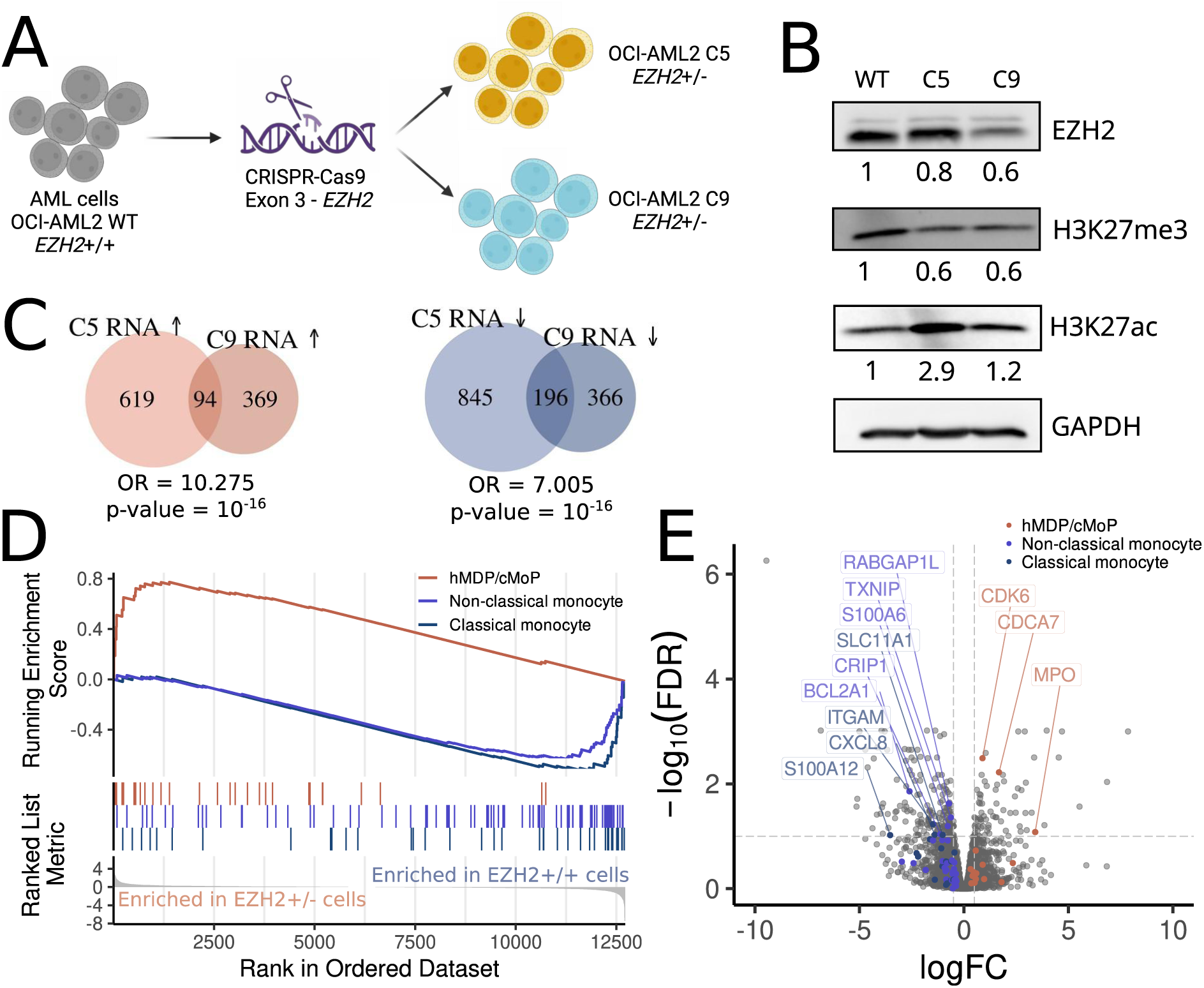
| PRC2 depletion leads to changes in gene expression related to cell differentiation. **A,** Two heterozygous EZH2 loss models (C5 and C9) were generated using CRISPR-Cas9 editing of the OCI-AML2 cell line. **B,** Immunoblotting of protein extracts from WT and EZH2-depleted cell lines using EZH2, H3K27me3 and H3K27ac antibodies, with GAPDH as a loading control. Quantification was performed using ImageJ. **C,** C5 and C9 differentially expressed genes from RNA-Seq. Red = upregulated; blue = downregulated; OR = Odds ratio for overlap; p-value was calculated using Fisher’s exact test. **D,** Most highly enriched gene sets from the atlas of human blood cells in OCI-AML2 PRC2-depleted (red) and PRC2-WT (blue). **E,** Core enrichment genes from gene sets enriched in panel D highlighted with blue (Monocytic genes) and red (hMDP/cMoP) in the volcano plot illustrating differentially enriched genes. Core enrichment genes with |Log2FC| > 0.5 and FDR < 10% are labelled.

We first assessed transcriptional differences in EZH2-deficient cells by RNA-sequencing (Supplementary Figure S1B). Differential expression analysis revealed that genes were up-and downregulated in similar proportions in both clones, with C5 having 619 genes upregulated and 845 genes downregulated, and C9 having 369 genes upregulated and 366 genes downregulated, compared with wild type (WT) transcription (Figure 1C, Supplementary Table S2). We found the overlap between clones to be significantly larger than expected by chance for both up- and down-regulated genes, with odds ratios (OR) of 10.275 (C5) and 7.005 (C9, Figure 1C), suggesting that a substantial proportion of the observed transcriptional differences were specifically linked to loss of EZH2 function.

We then performed gene set enrichment analysis (GSEA) to identify shared transcriptional signature changes between the two EZH2+/− clones. While global changes in pathway enrichment were limited (Supplementary Figure S1C), some specific differences were seen. As PRC2 is known to be critical for regulation of lineage-specific transcription in blood cell development, we were particularly interested in assessing expression changes in hematopoietic-related factors in more detail, in particular monocytic lineage genes that are most relevant for the OCI-AML2 myelomonocytic cell state. To do so, we leveraged publicly available scRNA-seq data from the Atlas of Human Blood cells (ABC) [26] to generate specific gene sets for the 32 ABC cell types (see Methods) for GSEA comparisons of EZH2 WT and EZH2+/− cells (Supplementary Figure S1D).

These analyses notably revealed a significant enrichment of immature human monocytic dendritic progenitors/common monocytic progenitors (hMDP/cMoP) genes upregulated in PRC2-depleted cells (NES = 2.18, FDR = 1.18×10^−4^) (Figure 1D, Supplementary Figure S1D). Conversely, we also observed a negative enrichment of differentiated monocyte program (i.e., classical monocytes: NES (Normalised enrichment score) = −1.92, FDR = 1.86×10^−4^ and non-classical monocytes and NES = −1.94, FDR = 7.57×10^−5^) (Figure 1D, Supplementary Figure S1D). PRC2-depleted cells show increased expression of hMDP/cMoP genes such as *MPO* and *CDCA7* (Figure 1E), while mature monocytic genes such as *CXCL8*, *S100A12,* and *ITGAM* (encoding CD11b) were downregulated (Figure 1E, Supplementary Table S2). Taken together, these results suggest that PRC2 depletion in this myelomonocytic cell line results in reorientation of transcriptional programs towards a more immature and less differentiated cell state (Supplementary Figure S1D).

### EZH2 depletion globally alters AML chromatin marks

To understand the epigenetic implications of depleting EZH2, we performed Cleavage Under Targets & Release Using Nuclease (CUT & RUN) [27] in WT and EZH2-deficient C5 and C9 cells to assess genome-wide deposition of PRC2-placed H3K27me3 and the transcriptionally activating mark H3K27ac. Principal Components Analysis (PCA) showed strong concordance between sample and histone modification replicates, allowing us to merge tracks for further analysis (Supplementary Figure S2A-B). We also confirmed strong concordance between H3K27me3 and H3K27ac signal in our samples and corresponding publicly available AML cell line data (Supplementary Figure S2C-E), suggesting that PRC2-mediated gene regulation is broadly conserved across different AML subtypes.

As expected with depletion of EZH2 K27 methyltransferase and PRC2 function, and in line with global H3K27me3 reductions (Figure 1B), there was ∼60% reduction in the total number of H3K27me3 peaks in our EZH2+/− models (124,100 peaks in WT vs 49,882 peaks in C5 and 46,165 in C9) (Figure 3A). Conversely, H3K27ac marks were increased (Supplementary Figure S2F). Overall, we observed that the H3K27me3 peaks conserved across WT and EZH2-depleted cells are enriched for promoter regions, whilst H3K27me3 peaks that were lost on EZH2 depletion are preferentially found in distal intergenic regions (Figure 2B). Similarly, H3K27ac peaks conserved across conditions are preferentially found at promoter sites and in gene introns. We also observed a slight enrichment for promoter sites among H3K27ac peaks that were lost in the clones (Supplementary Figure S2G).

**Figure 2.**
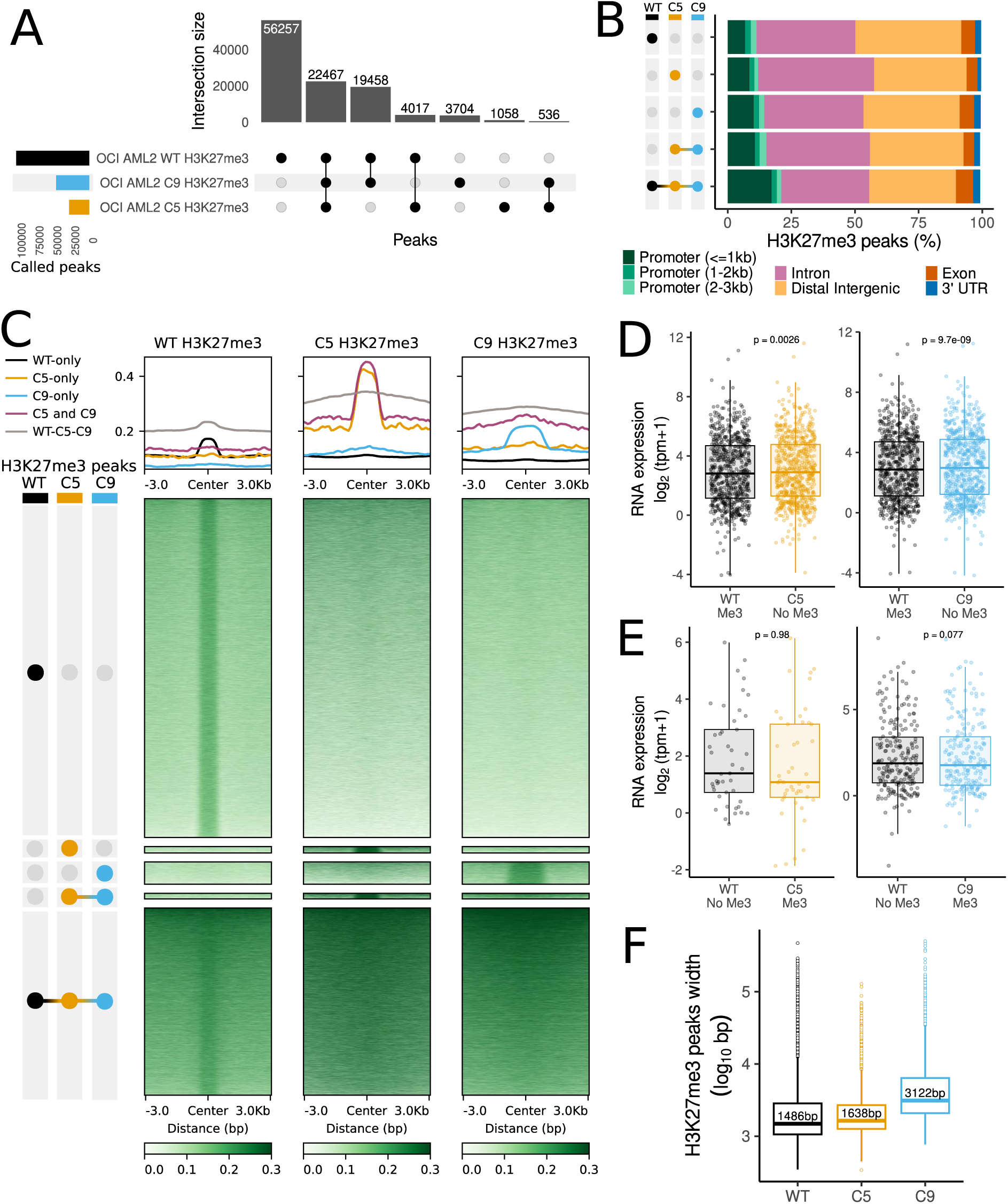
| PRC2 depletion leads to genome-wide decrease in H3K27me3. **A,** UpSet plot of H3K27me3 peaks called by SEACR in WT, C5 and C9 cells **B,** Annotation of H3K27me3 peaks called in WT, C5 and C9. **C,** Profile plots and heatmaps of H3K27me3 (green) signal at called H3K27me3 peaks stratified by sample in which the peaks were called. **D, E,** RNA expression in WT and C5 (orange) or C9 (blue) for genes that lose (**D**) or gain (**E**) H3K27me3 within the gene body or at the promoter upon PRC2 loss. P-values obtained upon comparisons using paired Wilcoxon tests. **F,** Width of H3K27me3 peaks from CUT&RUN in WT, C5 and C9.

**Figure 3.**
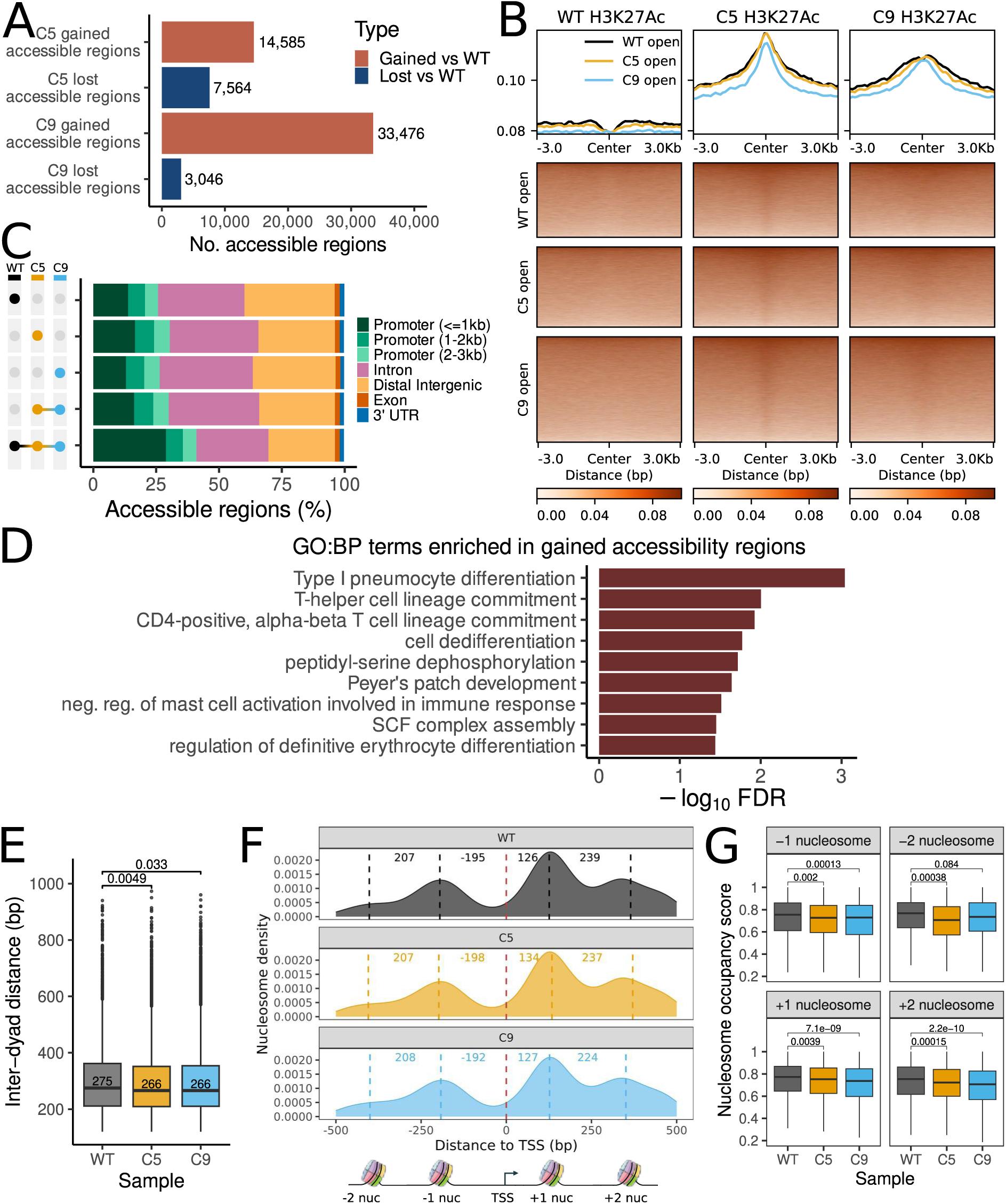
| Chromatin accessibility and transcriptional activity in PRC2-depleted AML cells. **A,** Chromatin accessibility profiles from ATAC-Seq summarised by comparing total number of accessible regions in C5 and C9 with total number of accessible regions in WT. **B,** Profile plots and heatmaps of H3K27ac (red) signal at called open chromatin peaks stratified by sample in which the peaks were called (WT-black, C5-orange, C9-cyan). **C,** Annotation by type of genomic regions for called accessible regions in the three conditions. **D,** GO enrichment of C5 and C9 accessible regions under an FDR < 5% and with a minimum of three genes annotated to the term. **E,** Inter-dyad distance for nucleosomes called using NucleoATAC near accessible regions at TSSs (Wilcoxon unpaired test p-value above brackets). **F,** Distribution of nucleosomes at genome-wide TSSs. Red dashed line = TSS. Black, yellow and blue lines = average nucleosome position. Values between dashed lines = average distance between TSS and nucleosome or between two neighbouring nucleosomes. **G,** Nucleosome occupancy score for −2, −1, +1 and +2 nucleosomes, respectively in WT, C5 and C9 (Wilcoxon unpaired test p-value above brackets).

Importantly, we found that genes that lost H3K27me3 at promoters or in the gene body had increased RNA expression (Figure 2D), linking EZH2 and PRC2 directly to transcriptional regulation of these loci. This included *CDCA7* from the hMDP/cMOP signature analysed above (Figure 1D-E, Supplementary Figure S1D, S2H). In contrast, we found no significant association between RNA expression and gain of H3K27me3 at promoter sites, with some genes becoming downregulated and others upregulated, despite deposition of repressive marks (Figure 2E). Interestingly, despite the observed global depletion of K27me3, we found that some regions gained stronger H3K27me3 signal in C5 and C9. These regions also became wider, suggesting more internucleosomal H3K27me3 spread in the EZH2+/− cells (Figure 2C, F).

We also found that H3K27ac did not show consistent links with transcriptional activity. H3K27ac changes were widespread, with 7,034 genes having gain of at least one peak and 8,957 having loss of at least one peak. Whilst 1,496 and 1,585 genes had increased H3K27ac levels in C5 and C9 respectively, only 9 genes with H3K27ac gain were transcriptionally upregulated in both clones (Supplemental Figure S2I). While some of these factors e.g., *CDCA7*, *LIN28B*, *CDK6* may be functionally relevant (see Figure 5), these results suggest that H3K27ac changes do not correlate directly with gene expression in our EZH2-low cells.

### Reduced PRC2 activity increases AML chromatin accessibility

Having assessed the effects of EZH2 reduction on histone modifications, we next wished to evaluate whether reduced PRC2 function and H3K27me3 depletion also affected chromatin accessibility. To analyse this on a genome-wide basis, we performed Assay for Transposase-Accessible Chromatin with high-throughput sequencing (ATAC-Seq) in our isogenic models. As PCA showed high similarity between experimental replicates (Supplementary Figure S3A), we combined cell line data for peak calling with HMMRATAC (see Methods).

Strikingly, we found large increases in the total number of peaks called in both C5 and C9, with 14,585 and 33,476 accessible peaks being gained respectively, in line with marked global increases in chromatin accessibility (Figure 3A, Supplementary Figure S3B). 10,601 of these i.e., the vast majority of new peaks in C5, were common to both clones (Figure 3A, Supplementary Figure S3B) and were enriched for H3K27ac (Figure 3B). This exceeds the number of lost accessible peaks by more than five-fold (Supplementary Figure S3B). Similar to CUT&RUN results, we found that regions with maintained accessibility across all conditions are enriched at promoter sites (Figure 3C, Supplementary Table S5).

### Open chromatin regions in PRC2-depleted cells are associated with development and cell differentiation

To interrogate functional differences between genes located in the increased and decreased accessible regions in EZH2+/− cells, we performed gene ontology (GO) enrichment analysis using Genomic Regions Enrichment of Annotations Tool (GREAT) [28,29]. We found no significant enrichment for any specific biological process in less accessible regions, suggesting that these changes do not markedly influence AML cell molecular phenotype. In contrast, there was significant enrichment of terms associated with hematopoietic development (e.g., T-helper cell lineage commitment) and broad development (e.g., cell dedifferentiation) in regions with increased accessibility in EZH2+/− cells (Figure 3D). This supports the hypothesis that PRC2 loss leads to opening of chromatin associated with hematopoietic differentiation and cell state. However, differential expression analysis showed that of the genes driving enriched GO terms in EZH2-deficient cells, only 2/40 are significantly upregulated (*AZU1*, *CDK6*, Supplementary Table S2, S6), suggesting that transcriptional relationships are incomplete.

To explore the links to gene expression further, we directly mapped fragments associated with open chromatin, i.e. nucleosome free regions (<100 bp), to transcriptional start sites (TSSs) of differentially expressed genes (see Methods). In both C5 and C9 we saw modest increases in TSS accessibility at upregulated genes (Supplementary Figure S3C) and slightly decreased TSS accessibility at downregulated genes (Supplementary Figure S3D). Visual inspection of the heatmaps suggested that most TSSs were already accessible in WT cells. In line with this, of 74 genes upregulated in EZH2+/− cells, only 19 were not expressed at all in EZH2+/+ cells, suggesting the majority of changes were quantitative in nature, rather than qualitative. We then analysed correlations between transcriptional and chromatin accessibility signals, and found that 134 and 117 genes had both increased promoter accessibility and increased RNA in C5 and C9, respectively (Supplementary Figure S3E, F). However, the vast majority of genes do not overlap, e.g., 1,517 genes have a more accessible promoter in C5 without showing significant RNA upregulation (Supplementary Figure S3E). Focussing on the overlap between the two clones, we identified 23 genes that increase both RNA and promoter accessibility in C5 and C9, which is significantly larger than expected by chance (OR = 4.469; Fisher’s exact test p-value = 10^-7^, Supplementary Figure S3G, H). Furthermore, we observe an overlap larger than expected by chance between C5 and C9 for genes that decrease in both RNA and accessibility (OR = 13.227; p-value = 5×10^−10^) (Supplementary Figure S3G, I).

We hypothesised that EZH2 depletion may not only affect chromatin accessibility, but also positioning of nucleosomes at TSSs [9]. To infer nucleosome positioning near open chromatin regions we used NucleoATAC [30]. We detected a slight increase in inter-dyad distance between nucleosomes (i.e., the length of DNA between neighbouring nucleosomes) in EZH2+/− cells, suggesting alteration of the nucleosome landscape (Figure 3E). We plotted the genome-wide nucleosome density at TSSs and observed a multi-modal distribution with four peaks, suggesting we can capture the −2, −1, +1 and +2 nucleosomes within ±500bp of the TSS (Figure 3F). We inferred the average positions of these nucleosomes by fitting a mixture model on each distribution (see Methods), and calculated the distance between each nucleosome and the TSS. Overall, we observed no difference in average nucleosome positions. However, nucleosome occupancy scores suggest a decrease in occupancy for all four inferred nucleosomes in EZH2+/− cells compared to EZH2+/+ (Figure 3G). This may be caused by a larger spread of nucleosomal fragments at these nucleosomes, meaning that nucleosomes are called with less confidence. This hypothesis is consistent with comparisons of nucleosome fuzziness scores, which are significantly higher in C5 at all nucleosome positions, and significantly higher in C9 at the −2 position (Supplementary Figure S3J).

Taken together our analysis suggests modest correlations between mRNA abundance and changes in chromatin accessibility, with a limited, but significant number of genes following a pattern of increased promoter accessibility that can be linked directly to increased transcription in EZH2+/− cells.

### Chromatin contact frequency impacts H3K27me3 retention

Our data suggest that the links between H3K27 methylation, chromatin accessibility and gene expression in AML are complex. We therefore speculated that additional factors influence transcription in EZH2+/− AML cells. As PcGs have been reported to control long range genomic interactions in other cellular contexts [7,17], we decided to investigate the role of PRC2 in 3D chromatin architecture in AML by performing Hi-C of the EZH2-depleted OCI-AML2 C9 cell line. We compared these results to publicly available OCI-AML2 WT Hi-C data [31]. OCI-AML2 has a highly disorganised genome, with many large-scale translocations and inversions that add unwanted noise in Hi-C data. Therefore, we performed Hi-C on an additional control cell line OCI-AML3 to control for both unwanted variation and batch effects.

After QC analyses, we calculated a matrix resolution of 15 kb for the Hi-C matrices of the three samples (i.e., more than 80% of 15 kb-sized bins contain more than 1,000 contacts) (Supplementary Figure S4A). Analysis of distance-decay curves revealed no significant differences across OCI-AML2 WT, OCI-AML2 C9, and OCI-AML3, indicating that there were no increases or decreases in loops of any specific size (Supplementary Figure S4B-C). As expected, a large proportion of loops (33%) called using Mustache [32] were present in all three samples, suggesting general conservation of global chromatin architecture (Figure 4A). Genome-wide pileups of these loops suggest a slight increase in looping intensity upon PRC2 depletion (Figure 4B). However, we do not see a clear change in the total number of loops, with roughly the same number of OCI-AML2 WT as C9 loops: 7,232 and 7,997 loops, respectively (Figure 4A).

**Figure 4.**
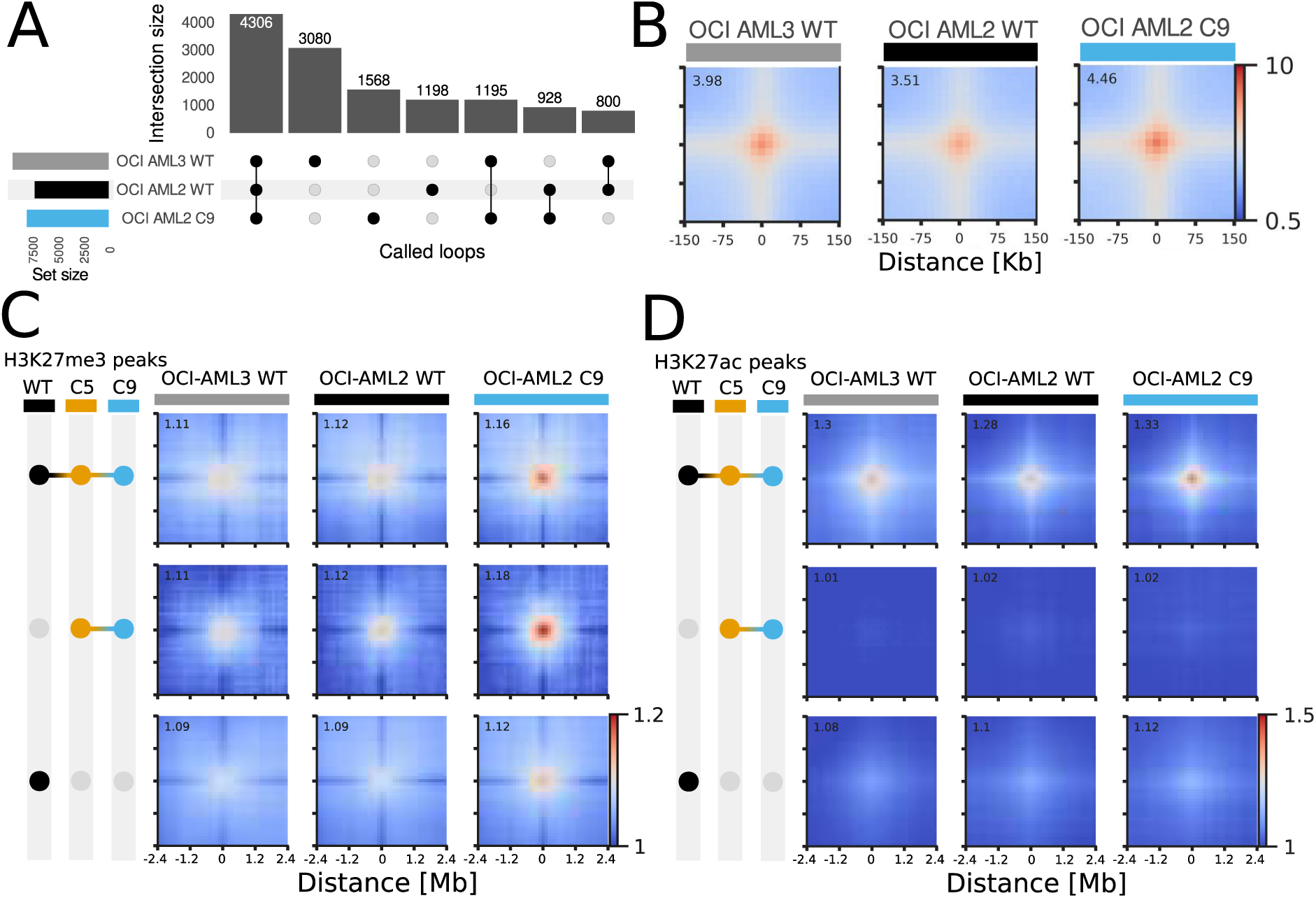
| The global chromatin looping landscape is altered upon PRC2 depletion. **A,** UpSet plot showing overlaps of loops called via Mustache in OCI-AML3, OCI-AML2 WT ([31]) and OCI-AML2 C9 Hi-C at 15kb resolution. **B,** Pileups of all called loops in the three conditions at 15kb resolution **C, D,** Pileups of contacts between domains of collapsed H3K27me3 (**C**) and H3K27ac (**D**) peaks (see Methods) and stratified by: WT-specific peaks (black),C5 and C9 union peaks (orange+cyan) and WT, C5 and C9 overlapping (black+orange+cyan) in the three cell lines at 120kb resolution.

We next investigated specific 3D chromatin changes at PRC2-bound loci, including the subset of H3K27me3 regions that are gained on EZH2 depletion (Figure 2F). As detailed earlier, these and other H3K7me3 peaks in EZH2+/− cells have a higher median width than those found in WT cells, suggesting both increased spreading and formation of broader H3K27me3 domains (Figure 3C, F). To gain insights into whether PRC2 binding influences genome architecture at these loci, we performed pile-up analysis of H3K27me3-decorated regions according to whether these were found only in WT, only in EZH2+/− clones, or were common to all conditions. As shown in Figure 4C, there were major differences in 3D chromatin interactions between these three categories. Firstly, the H3K27me3 regions that are lost upon EZH2 depletion do not seem to be involved in strong interactions. Secondly, the H3K27me3 regions that are maintained on EZH2 depletion show stronger DNA-DNA looping interactions in all cell lines, with the highest level of interactions in C9. Finally, on average, the newly deposited H3K27me3 marks are also enriched for 3D chromatin interactions, suggesting a role for PRC2 in maintenance of genome architecture at these sites. Additionally, we observed the same patterns of looping in publicly available Micro-C from the K562 cell line, suggesting that Polycomb-related loops are conserved across myeloid leukemia contexts (Supplementary Figure S4D).

The same analysis for H3K27ac peaks showed a similar pattern for conserved peaks, with those peaks present in all three conditions being heavily involved in looping, albeit with no observable differences between conditions (Figure 4D, Supplementary Figure 4D). In contrast to H3K27me3 peaks, H3K27ac peaks that were lost or gained on EZH2 depletion are not involved in looping (Figure 4D, Supplementary Figure 4D).

We also identified some short-range loops that were lost on EZH2 depletion which were linked to both H3K27me3 binding and gene expression, suggesting that these contacts are Polycomb-dependent (Supplementary Figure S5A). For example, H3K27me3 was increased at the boundaries of an ∼6Mb domain covering multiple long non-coding RNAs and *PCDH9* in EZH2-depleted cells, leading to formation of multiple loops within that domain and downregulation of *PCDH9* transcription (Supplementary Figure S5A-B).

### EZH2-dependent changes in chromatin architecture activate a LIN28B transcriptional signature and CDK6-related cell cycle disruption

Our results so far suggest that reduced EZH2 function has pleiotropic effects on gene regulation in AML cells. To assess whether any of these changes have biological significance, we next focussed on factors that are known to influence leukemia molecular phenotype.

We were particularly interested in the increased *LIN28B* expression that we detected in PRC2-depleted cells (logFC = 0.95, FDR = 0.04) (Supplementary Table S2). *LIN28B* is a fetal stem cell and lymphopoiesis marker that is rarely expressed in adult hematopoiesis [33], but which can be reactivated in several types of blood cancer. This can have clinical implications in some cases, e.g., *LIN28B* overexpression is associated with induction of a stemness transcriptional program and inferior patient outcomes in AML [34]. Because of this, and due to previously reported associations between *LIN28B* expression and EZH2 depletion in other contexts [18,35], we investigated further.

Strikingly, we found altered compartmentalisation near *LIN28B* at the 3D chromatin level, reflected by a “bowtie” shape that normally correlates with increased transcriptional activity (Figure 5A). These changes did not appear to be linked to H3K27me3 depletion, as this locus is already derepressed in OCI-AML2 according both to our measurements and Cancer Cell Line Encyclopaedia (CCLE) data (Supplementary Figure S5C). In contrast, the *LIN28B* locus is transcriptionally repressed and decorated with H3K27me3 in the MOLM13 AML cell line, as shown in comparisons in Figure 5A and Supplementary Figure S5C. We did however notice a small increase in H3K27ac across the *LIN28B* gene body in EZH2-depleted cells, which might be linked to the observed transcriptional changes (Figure 5B).

**Figure 5.**
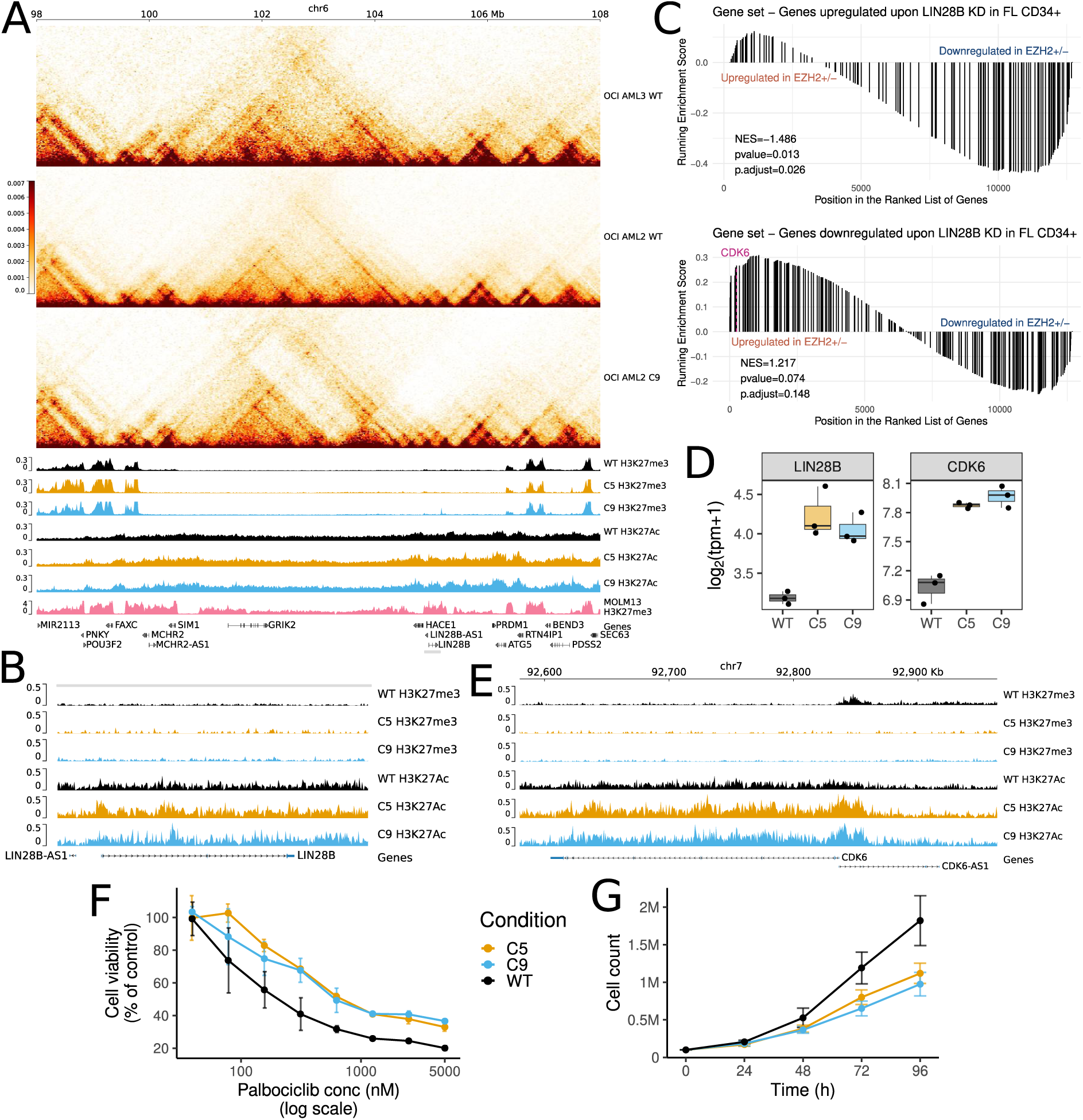
| Changes in chromatin organisation reveal a LIN28B signature contributing to drug resistance/cell cycle disruption through *CDK6* overexpression. **A,** Hi-C matrices for OCI-AML3, OCI-AML2 WT and OCI-AML2 C9 at the *LIN28B* locus at 30 kb resolution. H3K27me3 and H3K27ac tracks for WT (black), C5 (orange) and C9 (cyan) and H3K27me3 track from MOLM13 (pink) publicly available CUT&RUN data (see Methods). **B,** Zoomed in H3K27me3 and H3K27Ac tracks at the *LIN28B* gene. **C,** Gene set enrichment analysis on logFC ranked genes comparing EZH2+/− (C5 and C9 together) against EZH2+/+ cells. Gene sets tested were: genes up- and down-regulated upon LIN28B KD in foetal liver CD34+ cells (See Methods). D, *LIN28B* and *CDK6* mRNA expression in OCI-AML2 WT, C5 and C9. **E,** H3K27me3 and H3K27ac tracks for WT (black), C5 (orange) and C9 (cyan) **F,** Palbociclib treatment in OCI-AML2 WT, C5 and C9. Cell viability measured using Alamar Blue following 72 hr treatment with the indicated concentrations of Palbociclib. Data points represent the mean of N = 3 biological replicates, +/− SEM. **G,** Cell proliferation of WT, C5 and C9 cells, as measured by cell counting with Trypan Blue exclusion (N = 7 replicates +/− SEM).

To test whether *LIN28B* upregulation correlates with activation of a LIN28B-regulated transcriptional program, we used RNA-Seq data that was previously generated from human fetal liver (FL) CD34+ cells perturbed with *LIN28B* shRNA [36]. We derived two transcriptional signatures from these data, corresponding to genes upregulated or downregulated upon *LIN28B* knock-down (KD, see Methods). We saw that genes upregulated upon *LIN28B* KD in CD34+ FL cells were downregulated in our EZH2-depleted cell line model, consistent with LIN28B activation (NES = −1.486) (Figure 5C). In keeping with this, genes downregulated with LIN28B KD are upregulated in PRC2-depleted cells (NES = 1.217) (Figure 5C). This included LIN28B target genes involved in alternative lineage programs such as *ETS1*, *RAB7B* and *ZNF43 [37–39]*. Furthermore, and consistent with these results, we found *CDK6* to be highly downregulated in LIN28 KD CD34+ FL cells (logFC = - 1.53; FDR = 3×10^−39^) and upregulated in PRC2-depleted cells (logFC = 0.89, FDR = 0.003). These transcriptional changes correlated with chromatin marks at the *CDK6* promoter, as shown by decreased H3K27me3 and markedly increased H3K27ac in EZH2-depleted cells (Figure 5D-E). Taken together, these results suggest a potential association between PRC2 function and activation of a LIN28B transcriptional program that could influence leukemia biology.

Finally, we examined whether *CDK6* expression changes affect cellular phenotype. CDK6 is a cyclin-dependent kinase essential for the transition of cells from G1 to S phase and is essential for the progression and survival of many cancer types, leading to incorporation of CDK6 inhibition into cancer treatment strategies [40]. We therefore tested the effects of palbociclib, a CDK4/6 inhibitor which is currently approved for breast cancer treatment and is in clinical trials for AML and other cancers [41–43] in our cell line models. Strikingly, we found that PRC2-depleted cells were significantly less sensitive to palbociclib-induced cell death than WT cells (Figure 5F). Additionally, baseline cell proliferation was significantly lower in EZH2-depleted cells (Figure 5G). These results are in line with an elevated CDK6 activity in this setting, and correlate with reports of *CDK6* amplification promoting resistance to CDK6 inhibition in other cancer types [44].

## Discussion

Our results provide new mechanistic insights into the effects of PRC2 alteration in AML. We found that heterozygous loss of *EZH2* caused large increases in chromatin accessibility accompanied by elevated H3K27ac levels and decreased genome-wide H3K27me3. Gene expression changes included a notable activation of immature lineage transcriptional programs that may reflect reorientation towards a less differentiated cell state. Integration of 3D genome readouts further revealed a potential role for PRC2 in regulating a LIN28B-driven transcriptional program that appears to affect EZH2+/− cell phenotype through altered CDK6 activity.

While we found some correlations between chromatin accessibility and transcription, these relationships were incomplete. This may reflect the fact that some loci become ‘primed’ for activation, without necessarily initiating transcription. We do however see activation of immature monocyte lineage signatures and decreased expression of mature monocyte genes (e.g., *CD14*) that reflect a shift to a less differentiated transcriptional state in EZH2+/− cells. While we also detected changes at chromatin marks and accessibility at alternative lineage loci, in line with reports in other cellular contexts [45–47], it is likely that it is not possible to fully activate these programs in myelomonocytic OCI-AML2 cells, even if PRC2 repressive marks are removed. Despite some expected epigenetic and transcriptomic heterogeneity in the two EZH2+/− clones, we observed consistent similarities in chromatin accessibility near development-related genes and transcriptional activation of alternative lineage factors, including *LIN28B*.

We also showed that PRC2 depletion modifies the nucleosomal landscape in AML cells. In particular, we identified a decreased nucleosome occupancy score and increased nucleosome fuzziness near TSSs. This means that while nucleosomes remain, on average, at their normal positions, they may have more ‘wiggle room’ when PRC2 activity is reduced. A role for PRC2 in nucleosome positioning has not been fully clarified. Whilst we know that PRC1 depletion drastically changes the nucleosome landscape, SUZ12 depletion was reported to have no effect on nucleosome occupancy [8]. However, [9] showed that EZH2 depletion results in nucleosome repositioning at HOX gene promoters that are usually PcG-repressed.

While the role of PRC2 in maintaining 3D chromatin architecture in normal cells is well described [48–50], comparatively little is known about a corresponding function in leukemia. Hi-C analyses allowed us to detect that chromatin architecture is largely preserved in EZH2+/− AML cells. Strikingly, H3K27me3 was maintained in regions with high contact density, and H3K27me3 loss largely occurred in regions with less chromatin looping, suggesting that residual PRC2 function is important for maintaining genomic integrity at these sites. These results align with data from other contexts suggesting this is a conserved Polycomb function. For example, both PRC1 and PRC2 were reported to contribute to formation and maintenance of polycomb-associated domains in early mouse development [4]. In hematopoiesis, H3K27me3-mediated very long range interactions are essential in HSPCs, with pharmaceutical EZH2 inhibition causing loop relaxation and cell differentiation with loss of HSPC state [17]. Furthermore, Kraft et al. [7] show that polycomb-mediated 3D chromatin contacts are essential for H3K27me3 spreading, potentially explaining why ‘new’ H3K27me3 marks in EZH2+/− cells in our model are preferentially found at regions with high DNA-DNA contact frequency.

We further found that these chromatin architecture changes may influence leukemia phenotype, thereby providing biological relevance. In this case, EZH2 depletion led to genome compartmentalisation changes at the *LIN28B* locus. While this gene is already expressed in OCI-AML2, increased transcription in EZH2+/− cells was sufficient to activate a LIN28B-driven transcriptional signature, as previously described in other blood cell models [15,18]. LIN28B is an RNA-binding protein that is rarely expressed in normal adult cells, but which is active in > 20 cancer types, including leukemias [51]. LIN28B mainly acts as an inhibitor of tumour suppressive let-7 micro-RNAs that normally repress oncogenes and cell cycle drivers such as *MYC*, *RAS,* and *HMGA2* [52,53][54][52,53].

Part of this LIN28B-driven signature is *CDK6*, which shows transcriptional upregulation and loss of promoter H3K27me3 in EZH2+/− cells. CDK6 mediates G1/S transition and can be overexpressed in acute leukemias [55]. *CDK6* is often upregulated in *KMT2A*-rearranged (*KMT2A*r) AML and is a direct target of KMT2A fusion proteins [56], leading to its being proposed as a therapeutic target for these cases. Interestingly, treatment with the CDK6 inhibitor palbociclib has been shown to inhibit *KMT2A*r cell growth and to affect expression of myeloid differentiation genes [56]. In the case of our experimental model, EZH2 depletion caused reduced cell proliferation and corresponding decreased sensitivity to palbociclib, identifying potential issues with implementing this treatment in PRC2-altered AML.

Several key questions remain to be addressed before the role of EZH2 in leukemia transcriptional regulation can be fully understood. For example, knowledge of the dynamics by which PcGs suppress alternative lineages in blood cells, similarly to what has been recently described during endodermal differentiation [47], is currently limited. Single-cell studies investigating transcription, chromatin accessibility and response to perturbation may be able to disentangle the role of EZH2 in lineage priming in this context. Furthermore, intra- and inter-leukemia heterogeneity may be driven by the presence of other oncogenic alterations, such as the *KMT2A* rearrangement present in our cell line model, or additional driver mutations. There is a need therefore to study how different genetic contexts modify PRC2 epigenetic activity, and how the effects of different oncogenic drivers functionally interact to modify leukemia transcription and biology.

## Conclusions

This study provides a comprehensive characterization of transcriptional changes caused by PRC2 depletion in AML cells. At the epigenome level, we found that heterozygous *EZH2* loss results in marked decrease of H3K27me3, accompanied by increased H3K27ac and significantly increased chromatin accessibility. Strikingly, we found that H3K27me3 is maintained at genomic regions with high contact frequency. These epigenomic changes could be linked to transcriptional activation of immature and alternative lineage programs, including an oncofetal *LIN28B*-driven program that was associated with *CDK6* upregulation and resistance to CDK6 inhibitor palbociclib. Taken together, these findings provide new insights into the mechanisms by which altered PRC2 function impacts transcriptional regulation, molecular phenotype, and treatment responsiveness of leukemia cells.

## Methods

### CRISPR/Cas9 gene editing of AML cell lines

#### Electroporation

Two guide-RNAs (gRNA, protospacer adjacent motif or PAM sequence underlined, #1: CGGAAATCTTAAACCAAGA<u>ATG</u>, #2: ACCAAGAATGGAAACAGCG<u>AAG</u>), were designed to specifically direct the Cas9 endonuclease to the third exon of our gene of interest, *EZH2* (ENST00000460911.5; chr7:148,543,562-148,543,689 Reverse strand, GRCh37/hg19). gRNAs were purchased from Integrated DNA Technologies.

To deliver both the Cas9 enzyme and *EZH2*-targeted gRNAs into the OCI-AML2 cell line (gift from Mills Laboratory, Queen’s University Belfast), we utilised the Alt-R CRISPR-Cas9 System from Integrated DNA Technologies. This approach allowed incorporation of both components into the cells, following electroporation, as a ribonucleoprotein (RNP) complex. Electroporation was performed using the Cell Line Nucleofector™ Kit V (Lonza, cat. no. VCA-1003) coupled with the Amaxa® Nucleofector® System (Lonza, Nucleofector 2b) according to the manufacturer’s instructions. Lyophilised gRNAs were reconstituted in IDTE buffer at a final concentration of 200 µM. To achieve the recommended total amount of guide RNAs (100 µM) using 2 separate gRNAs, a 1:1 mixture was made in which each gRNA was at a concentration of 50 µM. The gRNA mixture was combined with 104 pmol of recombinant Cas9 protein and incubated for 20 minutes at room temperature to allow formation of the RNP complex. Cells were rinsed with PBS, counted using trypan blue exclusion, centrifuged, and resuspended in 60 µL Nucleofector^TM^ solution at a density of 16,500 cells/µL. 15 µL of the RNP complex solution was added to the cells, as well as 3 µL of the Alt-R Cas9 Electroporation Enhancer solution (previously resuspended in IDTE buffer at a concentration of 100 µM). Nucleofection was performed using the Amaxa Nucleofector 2b device, using the electroporation settings outlined above. Following electroporation, cells were transferred into 500 µL of pre-warmed media per well of a 24 well plate and stored at 37°C, 5% CO_2_.

#### Selection of single-cell clones

48 hours post-electroporation, the bulk transfected cells were isolated into single cells to allow growth of single cell colonies. For this, cells were counted using trypan blue exclusion and diluted to a concentration of 5 cells/mL. The cell suspension was then distributed in 96 well plates, adding 100 µL of cell suspension in each well. Using an initial concentration of 5 cells/mL per well (0.5 cells per well) minimised the probability of seeding more than one cell per well.

Cells were maintained in RPMI-1640 media supplemented with 20% fetal bovine serum (FBS) and 2 mM L-Glutamine. Once the cells reached confluence, they were successively transferred to 24-, 12-, and then 6-well plates. When the cell number was sufficient, proteins were extracted, and the lysates were run on a 10% acrylamide gel and subjected to immunoblotting to assess EZH2 protein levels. DNA was then extracted from clones showing reduced EZH2 levels, using the DNeasy® Blood and Tissue Kit (Qiagen, Cat 69504). Genomic DNA was used to amplify Exon 3 of *EZH2* via PCR. Amplicons were then subjected to direct (Sanger) sequencing to detect potential insertions or deletions (indels) responsible for reduced EZH2 levels. Cell line identity was authenticated by single nucleotide polymorphism profiling using a commercial service (Eurofins) and all cell lines used in this study were tested regularly (at least every 3 months) for mycoplasma using Lonza’s MycoAlert® Mycoplasma Detection Kit (LT07-710) according to the manufacturer’s instructions.

#### Identification of indels induced by CRISPR/Cas9 editing

Sanger sequencing results for *EZH2*-WT and candidate *EZH2*-knock-out (KO) cells were analysed using the ICE (Inference of CRISPR Edits) CRISPR Analysis Tool (available at https://ice.synthego.com), which integrates gRNA sequences and DNA sequence electropherogram files (.ab1 format). The ICE tool provides a knock-out (KO) score, which is the proportion of cells that have a frameshift or an indel greater than 21 base-pairs in length. Identification of a frameshift or indel using this tool allows for the assessment of functional consequences to the gene and its protein product.

### Immunoblotting

#### Cell lysis

Cells were harvested (centrifugation at 350 xg) and pelleted. Protein lysates were obtained from a minimum of 1×10^6^ cells lysed in a home-made lysis buffer (1M Tris pH7.5, 5M NaCl, and 0.5% (v/v) NP40, H_2_O, filtered), supplemented with protease and phosphatase inhibitors (1 Tablet of each/10mL, COmplete Mini Protease Inhibitor Cocktail Tablets – Roche 11836153001; PhosSTOP Phosphatase Inhibitor Cocktail Tablets – Roche 4906837001). Briefly, cell pellets were resuspended with a minimum of 120 μL of lysis buffer. Tubes were briefly vortexed and left on ice for 15 minutes before being vortexed again and centrifuged for 10 minutes at 18,000 xg at 4°C. The supernatant (i.e., cell lysate) was collected and was either used immediately or stored at −20°C. To assess the protein concentration in cell lysates, the Pierce^TM^ BCA Protein Assay Kit (Thermo Scientific - cat number: 23227) was used according to the manufacturer’s instructions..

#### Western blotting

10 to 20 μg of protein/sample were mixed (4:1 v/v) with a mix of NuPAGE LDS Sample Buffer (1:4) containing 1M dithiothreitol (DTT), incubated for 5 minutes at 95 °C and then resolved on 10% acrylamide gels via electrophoresis. Next, separated proteins were transferred to a methanol-activated PVDF membrane using a semi-dry transfer method (transfer buffer: 10% 20X Tris-glycine and 20% methanol in MiliQ water). The membrane was then blocked with 5% (w/v) skimmed milk powder dissolved in 1x TBS-T (Tris Buffered Saline-Tween) shaking for 1 hour at room temperature.

The membrane was then incubated overnight with primary antibody at 4 °C (Supplementary Table S1). After washing with 1x TBS-T, the membrane was incubated for 1 hour with an HRP-conjugated secondary antibody. An in-house enhanced chemiluminescence (ECL, Solution 1: 1M Tris/HCl pH8.5, p-coumaric acid, luminol and H_2_O; Solution 2: 1M Tris/HCl pH8.5, 30% Hydrogen peroxide (H_2_O_2_), and H_2_O) solution (1:1) was applied to the membrane for protein detection using the Advanced Molecular Vision Chemi Image Unit of the ChemoStar Imager (INTAS Science Imaging Instruments GmbH).

### *In vitro* proliferation assay

100,000 cells were seeded in 1 mL of complete media in 24-well plates and incubated at 37℃ in 5% O_2_ in a humidified incubator. Viable cells were manually counted using a haemocytometer following trypan blue staining at 24, 48, 72 and 96 hours.

### RNA-Seq

#### Library generation

RNA was extracted from 3-5 million cells using the RNAeasy kit (Qiagen, 74104). Three independent RNA extractions were performed at serial passages for all cell lines to provide biological replicates. RNA quality was assessed using the Agilent 2100 Bioanalyzer prior to submission for sequencing as a service provided by Novogene UK. Before library preparation, samples were first enriched using oligo(dT) beads, mRNA was then randomly fragmented (average length of reads: 150 bp) before cDNA synthesis by reverse transcriptase. Library preparation was then carried out and library QC analysis was performed prior to paired-end sequencing at a depth of 30 million reads, resulting in .fastq files for processing.

#### Sequencing data processing

We pseudoaligned the reads against the GRCh38 (hg38) human reference genome using kallisto 0.46.1 [57] to obtain gene-level read counts. Quality Control (QC) was performed using FastQC 0.11.9 and MultiQC 11.9. Subsequent analyses were performed using R v4.4.0 (https://www.R-project.org/) and Bioconductor 3.19 [58].

#### Differential expression analysis

We normalised the read counts to log2 transcripts per million (TPM) and Trimmed Means of M values (TMM) using the edgeR package v4.2.0 [59]. Then, we filtered out lowly expressed genes (<1 counts per million (CPM) in more than three samples). We performed variance-stabilisation with the voom function from limma v3.60.3 [60]. To identify differentially expressed genes (DEGs) we used a limma linear model and we adjusted p-values for multiple testing using the Benjamini-Hocheberg correction. We identified significantly up- and down-regulated genes based on a false discovery rate (FDR) of 10% and |logFC| ≥ 0.5. Overlaps between differentially expressed genes were plotted using the VennDiagram v1.6.0 R package. Odds ratios and p-values for overlaps were calculated using the fisher.test function from the stats package.

#### GSEA

We performed gene set enrichment analysis (GSEA) through the clusterProfiler package v4.12.0 [61] using the Hallmarks (H) gene sets from MSigDB [62]. We identified significantly enriched gene sets using an FDR of 5%. We performed GSEA using gene signature from the Atlas of Human Blood cells [26]. We considered genes most highly expressed in each cell type as a separate transcriptional signature and identified significantly enriched cell type transcriptional signatures under an FDR of 10%.

### CUT&RUN

We performed Cleavage Under Targets & Release Using Nuclease (CUT&RUN) on WT, C5 and C9 for H3K27me3 and H3K27ac in duplicate in addition to one sample IgG per condition.

#### Binding cells to Concanavalin A beads and primary antibody

500,000 OCI-AML2 WT, C5, and C9 cells per CUT&RUN condition and for input condition were harvested at 700 xg for 3 minutes. To each sample, 37% formaldehyde to a final concentration of 0,1% was added, and was quenched using 1/20^th^ volume 2.5M Glycine (final concentration 125 mM Glycine) after 1 minute. Cells were then spun for 3 minutes at 600 xg and then washed once in PBS. Pelleted cells were then resuspended in 100 μL/sample in cold Nuclear Extraction Buffer (supplemented with spermidine and phosphatase and protease inhibitors). After a 10-minute incubation on ice, samples were spun for 3 minutes at 600xg and input samples were then stored on ice. The resulting nuclei were resuspended in 100 μL/sample in cold Nuclear Extraction Buffer and aliquoted to each 8-strip tube containing 10 μL/sample previously-activated Concanavalin A beads (Cell Signaling Technology, Cat. 93569S) and placed on the rotator to incubate for 10 minutes at room temperature. Beads were isolated using a magnetic rack and resuspended in a 200 μL buffer containing the antibody of interest (Supplementary Table S1) in a 1:50 (v/v) final concentration. These tubes were rotated overnight at 4°C.

#### NGS library preparation

NGS libraries for Illumina were prepared using the NEBNext UltraExpress kit (NEB, Cat. E3325S) following the manufacturer’s instructions. 25 ng purified DNA were end prepped using the provided enzyme mix in a thermocycler (20°C for 30 minutes then 65°C for 30 minutes). The NEB adaptor (P*-GATCGGAAGAGCACACGTCTGAACTCCAGTC-U-ACACTCTTTCCCTACACG CTCTTCCGATC*T) was then ligated to the end prep reaction mixture using the NEBNext Ligation Master mix and incubated for 15 minutes at 20°C. Following this incubation, 2 μL USER enzyme was added to each tube and allowed to incubate for a further 15 minutes at 37°C.

Adaptor-ligated libraries were cleaned using the Beckman Coulter Agencourt AMPure XP beads (Thermo Scientific, Cat. 10136224). A volume of 1.1x resuspended beads were added to the adaptor-ligated reaction. The solution was incubated on the benchtop for 5 minutes, then placed on a magnetic rack. After the supernatant was discarded, beads were washed twice with 200 μL 85% ethanol. DNA from the beads was ultimately eluted using 17 μL of 0.1X TE buffer. The adaptor-ligated cleaned up product was then PCR amplified using the provided NEBNext MSTC High Yield Master Mix and indexed using a combination of the i7 and i5 Illumina primers. Following one cycle of initial denaturation (98°C for 45 seconds), the sample underwent further denaturation (98°C for 15 seconds) and annealing/extension (60°C for 10 seconds) for 14 cycles. Final extension was then performed for one cycle at 72°C for 1 minute. Lastly, PCR-amplified libraries were cleaned using the Beckman Coulter Agencourt AMPure XP beads. A volume of 1.1x resuspended beads were added to the PCR reaction. The solution was incubated on the benchtop for 5 minutes, then placed on a magnetic rack. After the supernatant was discarded, the beads were washed twice with 200 μL 80% ethanol. DNA from the beads was ultimately eluted using 15 μL of 0.1X TE buffer. DNA concentration was measured using the Qubit dsDNA High-Sensitivity kit (Invitrogen, Cat. Q32851), while the size distribution of the libraries was assessed using the Agilent Bioanalyzer High Sensitivity DNA chip following manufacturer’s instructions.

#### Deriving an Input sample for RT-qPCR

The input samples stored on ice were resuspended in 200 μL cold 0.9% sonication buffer (supplemented with phosphatase and protease inhibitors). The input samples were then sonicated using a probe sonicator (Benchmark Pulse 150 Ultrasonic Homogeniser) for 19 cycles of 10 seconds sonication and 10 seconds break. Lastly, the DNA was centrifuged at 20.000xg for 15 minutes (4°C) and 180 μL supernatant stored at −20°C until further use.

#### Binding of pAG-MNase and targeted chromatin digestion

Beads were isolated using a magnetic rack following antibody binding and resuspended in 250 μL Digitonin-Wash buffer. pAG-MNase was then added to the tube in a final concentration of 0.9 ng/μL and incubated for 10 minutes at room temperature. Following incubation, the beads were isolated using a magnetic rack and resuspended in a 250 μL Digitonin-Wash buffer. Following two washes, beads were resuspended in 50 μL cold Digitonin Buffer. While incubating on ice, 1 μL 100 mM CaCl2 was added to each tube to activate the pAG-MNase. After two hours incubation at 4°C, 33 μL STOP-buffer was added to each tube and incubated at 37°C for 10 minutes to release soluble chromatin fragments. After incubation, 0.68 μL 10% SDS and 1 μL proteinase K (20μg/μL) was added to the 75 μL Cut&Run sample and then incubated overnight at 55°C. Input samples were treated with 3.46 μL RNase A (10mg/mL) and incubated at 37°C for 2 hours. Following this 1.62 μL 10% SDS and 1.73 μL proteinase K (20μg/μL) was added to the 180 μL sonicated input chromatin and incubated at 55°C overnight.

#### DNA isolation and purification

Following overnight incubation, 2 μL were taken from the input sample tubes and diluted in 73 μL 0.9% sonication buffer. Soluble chromatin fragment DNA and DNA from diluted input samples were isolated using the MinElute PCR Purification kit (Qiagen, Cat. 28004) following the manufacturer’s instructions. In short, 5 volumes of Buffer PB were added to 1 volume of digested or input samples, placed on the provided MinElute column, centrifuged at 17,900 xg for 1 minute, then washed with 750 μL Buffer PE, centrifuged twice for 1 minute each at 17,900 xg, then eluted in 21 μL Elution buffer. Input DNA was diluted in 100 μL Elution buffer. Using the Qubit dsDNA High-Sensitivity kit (Invitrogen, Cat. Q32851), 1 μL DNA was quantified on the Qubit fluorometer.

#### RT-qPCR

To validate successful antibody binding and chromatin release. Primers were designed around the transcriptional start site of *GAPDH*, the negative control, and the promoter region of *HEY2*, the positive control. 2 μL isolated DNA was pipetted in duplicates for each target gene in a 384-well plate. PCR-grade water was used as a non-target control (NTC). To each well containing DNA, 8 μL master mix per sample (containing 5 μL SYBR safe PCR mix (PowerUp SYBR Green Master Mix, ThermoFisher), 2.5 μL PCR-grade water, and 0.25 μL 10 μM forward and reverse primers) was added. The plate was then incubated in a QuantStudio 7Flex qPCR system with the following programme: in the hold stage, samples were incubated at 50°C for 2 minutes and at 95°C for 20 minutes (1.6°C/s temperature increments). During the PCR stage, samples were incubated at °C for 15 seconds and 60°C for 1 minute for a total of 40 cycles, while in the melt curve stage, samples were kept at 95°C for 15 seconds, 60°C for 1 minute and 95°C for 15 seconds to ultimately achieve dissociation. To assess whether there is an enrichment in the target of interest, comparative Ct (ΔΔCt = ΔCt _antibody of interest_ – ΔCt_input_) was calculated. From this, the fold-change was calculated (fold-change = 2^(-ΔΔCt)^). A fold-change of over 1 over the IgG control was used to determine successful antibody binding to DNA in the positive control.

#### NGS data processing

QC of the unaligned reads was performed using FastQC 0.11.9 and MultiQC 11.9. We performed adapter (TrueSeq adapters) removal with Trimmomatic v0.39 on the paired reads with settings as recommended by the 4DN project [63]. We aligned the reads against the human reference genome hg38 using bowtie2 v2.3.5.1 --end-to-end with the following parameters: inclusion of dovetailed reads (--dovetail), only concordant (--no-discordant), paired reads only (--no-mixed), and with lengths between 10 and 700 bp (-I 10 -X 700). We marked and removed duplicates and multi mappers using Picard v3.1.1 and samtools v1.18 (-F 1280 parameter). For visualisation purposes we created .bigwig files using bamCoverage using --binSize 30 --smoothLength 60 --normalizeUsing CPM -- effectiveGenomeSize 2913022398 --extendReads. We used these .bigwig files for generating deeptools heatmaps and profile plots.

#### Peak calling and annotation

We performed peak calling following the recommendations from SEACR [64]. Specifically, we used the .bam files to obtain .bedgraph files through bedtools genomecov and called stringent peaks for each condition using SEACR v1.3 normalised to IgG. For heatmaps and profile plots over .bigwig files, we used the computeMatrix and plotHeatmap functions from deeptools v3.5.4 [65]. We found unique and overlapping peaks across the two conditions and two histone marks using the vennCount function with maxgap=1000bp from the hicVennDiagram v1.2.0 R package. We annotated the called CUT&RUN peaks with ChIPseeker v1.5.1 with a TSS region spanning between −3kb and +3kb and the parameter overlap = “all” to limit bias towards TSS annotation [66].

Figures containing tracks of ATAC-Seq, CUT&RUN or Hi-C were obtained using pyGenomeTracks v3.9 [67].

### ATAC-Seq

100,000 cells from each of the conditions (OCI-AML2 WT, C5 and C9) were harvested by centrifugation at 500 xg at 4°C for 5 minutes and subsequently lysed in an ice-cold ATAC Lysis buffer.

#### NGS library preparation

Assay for Transposase-Accessible Chromatin with sequencing (ATAC-Seq) was performed using an Active Motif kit (Cat. no. 53150) according to the manufacturer’s instructions. The eluted DNA was subsequently amplified using a combination of indexed primers i7 and i5 (25 μM) in a thermocycler with the heated lid on with the following steps: 72°C for 5 minutes, 98°C for 30 seconds, followed by 10 cycles of 98°C for 10 seconds, 63°C for 30 seconds, and 72°C for 1 minute. Amplified DNA was size-selected using 1.2x SPRI beads. These were then washed using 180 μL 80% ethanol twice and DNA was ultimately eluted using 25 μL elution buffer. The complexity of the library was assessed using the Agilent Bioanalyzer and DNA was quantified using the High-Sensitivity DNA Qubit kit (Cat no. Q32851, ThermoFisher Scientific)

#### NGS data processing

We applied the nf-core/atacseq v2.1.2 pipeline to perform initial QC, adapter trimming, duplicate removal and alignment of reads into .bam files. We used the peak calls from the pipeline to construct a PCA in order to check for similarity between replicates. Based on this output, we concluded that the replicates were similar (Supplementary Figure S3A) and used the merged .bam files to apply custom pipelines as described below.

#### Peak calling

We performed peak calling using HMMRATAC v1.2.10 using default settings [68] and excluding the hg38 v2 blacklist from https://github.com/Boyle-Lab/Blacklist [69].

#### Peak overlaps and annotation

We found unique and overlapping peaks between the three conditions using the vennCount function with maxgap=50bp from the hicVennDiagram v1.2.0 R package. We annotated the called peaks with ChIPseeker v1.5.1 with a TSS region spanning between −3kb and +3kb and the parameter overlap = “all” to limit bias towards TSS annotation.

#### Heatmaps

Using alignmentSieve from deeptools v3.5.4, we separated the ATAC-seq data for each condition into two groups based on fragment size: nucleosome-free regions (NFRs) <100bp and mononucleosomal fragments between 180 and 250bp. We generated bigwig files from .bam files using bamCoverage and the following parameters: --binSize 1 --normalizeUsing RPGC -- effectiveGenomeSize 2913022398. For heatmaps and profile plots over .bigwig files, we used the computeMatrix and plotHeatmap functions from deeptools v3.5.4. All heatmaps for ATAC-seq were generated using NFR regions.

#### Enrichment analysis

We performed genomic regions enrichment on the overlapping C5 and C9 open regions, using all called peaks as background. For this analysis we used the web-based tool GREAT with default parameters: http://great.stanford.edu/ [28]

#### Nucleosome positioning inference

We inferred nucleosome positions around NFRs using NucleoATAC 0.3.4 with default settings [30]. For each nucleosome inferred, NucleoATAC provides an occupancy score and a fuzziness score. Median inter-dyad distances were compared across the three conditions using Wilcoxon unpaired tests. We stratified the inferred nucleosomes within ±500bp of TSSs into - 2, −1, +1 and +2 nucleosomes by fitting a mixture model with four components to the inferred nucleosome positions using the mixtools R package and allocating each nucleosome to one of the preset positions based on its likelihood. After this, we used the average position of each nucleosome type (−2, −1, +1 and +2) to calculate the average distance from TSS. Median nucleosome occupancy scores and nucleosome fuzziness were compared across the three conditions using Wilcoxon unpaired tests.

### Hi-C

#### Library preparation

All steps for the generation of Hi-C samples for sequencing (sample processing and library preparation) were performed as a service by Active Motif using the Arima-HiC Kit. 10^7^ cells per cell line were washed in 1x PBS, pelleted and stored at −80°C prior to submission. The Active Motif Hi-C workflow used the Arima-HiC Kit (Arima Genomics) for performing chromatin conformation capture (crosslinking, digestion, biotinylation, ligation and fragmentation). The Active Motif workflow for library preparation used the KAPA Library Amplification Kit (Roche), according to manufacturer’s instructions. Hi-C libraries were diluted to 20 nM and submitted for sequencing to Novogene. Sequencing of the pre-made libraries was performed, generating 300 million reads (paired-end) per sample.

#### NGS data processing

We performed QC using FastQC v0.12.1 and aggregated the outputs using MultiQC. We aligned the raw reads against the reference human genome hg38 using BWA-MEM2 v2.2.1 [70]. We then created pair files using pairtools v1.0.3 with a minimum mapping quality (mapq) of 3 and the parameter “--report-position outer”, as recommended on the Pairtools Github https://github.com/open2c/pairtools [71]. We used pairtools to further split and sort the paired reads. We then used the FAN-C command “fanc pairs” with the “-restriction_enzyme” argument to filter for ligation errors [72]. For OCI-AML2 we used the DpnII restriction fragments as input, as DpnII was used to generate the data, as stated by [71]. For the Hi-C data generated by us (OCI-AML3 and OCI-AML2 C9) we used the Arima-specific list of restriction fragments (i.e., Arima uses both HinfI and DpnII). After investigating ligation error statistics, we filtered the paired reads using fanc pairs with the following parameters: “--filter-unmappable --filter-multimapping -- filter-inward 5000 --filter-outward 5000 --filter-self-ligations --filter-pcr-duplicates 1”. We used “fanc hic” to create a .hic matrix from the .pairs file and then applied ICE (iterative correction and eigenvector decomposition) normalisation.

#### Cooler files generation

To calculate the resolution of each .hic matrix, we used “fanc resolution”. More precisely, for bin sizes of 5kb, 10kb, 12kb, 15kb, 20kb, 50kb, 100kb, in each matrix, we calculated the percentage of bins with >1000 contacts. We set a threshold at 80% of bins having >1000 contacts and concluded that the optimal resolution for the .hic maps was 15kb. We created .hic maps at 7.5kb resolution for visual inspection purposes and used fanc to-cooler with the parameters “--uncorrected” and “--no-multi” to generate a .cool file. Further, we used the zoomify function from cooler 0.9.3 to generate an .mcool file with decreased resolutions (15kb, 30kb, 60kb, 120kb, 240kb etc.). We used the “--balance” argument for normalisation and “--mad-max 0 --max-iters 1000” as suggested on https://github.com/open2c/cooler.

#### Distance decay

We calculated distance decay for each condition using fanc expected. The distance-decay curve allows us to visualise the frequency of contacts as a function of genomic distance. The derivative of the distance-decay curve allows us to better visualise any change in directionality of the curve.

#### Loop calling

We performed loop calling at 15kb resolution using Mustache v.1.3.2 with σ_0_=1.6 [32]. For determining condition-specific and overlapping loops, we used the hicVennDiagram v1.2.0 R package with a maxgap=20kb.

#### Pile-ups

We performed pile-ups of Mustache loop calls using coolpup.py v1.1.0 [73] at 15kb resolution with 150kb flanking in each direction from the loop centre. Similarly, we performed pile ups on interactions between H3K27me3 regions. Due to the maximum resolution of the Hi-C data being 15kb, we decided to merge H3K27me3 loops into domains using bedtools, as follows: if the distance between two loops is <15kb, they are collapsed within a domain (bedtools merge -d 15000). Then we performed pileups at 15kb, 30kb, 60kb, 120kb and 240kb with 20x resolution as flanking. We observed that the domains were most likely larger than we expected, hence the 120kb resolution pileups looked the best (i.e, there was a visible central enrichment smaller than the size of the pileup window).

For visualisation of Hi-C maps we used HiGlass [74] and pyGenomeTracks [67].

### *In vitro* efficacy of palbociclib against AML cell lines

Cells were plated at a density of 8,000 cells/well in 96-well U-bottom plates. Cells were equilibrated overnight at 37 °C, 5% CO2 prior to drug treatment. Palbociclib (MedChemExpress) was serially diluted (1:2) in culture media and dilutions added in triplicate wells, with control wells containing 0.1% DMSO. Following 72 h drug exposure, cell viability was assessed by mitochondrial activity assay (Resazurin cell viability assay). Resazurin solution (0.6 mmol/L Resazurin, 0.07 mmol/L Methylene Blue, 1 mmol/L potassium hexacyanoferrate (III), 1 mmol/L potassium hexacyanoferrate (II) trihydrate) was added to all wells and plates incubated for afurther 4 h at 37 °C, 5% CO2. Luminescence was read using a SpectraMax M3 plate reader (Molecular Devices) with excitation at 560 nm and emission at 590 nm. Cell viability was calculated as the percentage of vehicle-treated controls.

### Publicly available data

Gene signatures based on scRNA-Seq from the Atlas of Blood Cells were downloaded from http://scrna.sklehabc.com/ on 22nd of July 2025. These data were published by Xie et al. [26]. H3K27me3 CUT&RUN signal in .bigwig format and peak calls in .broadPeak format from the MOLM13 cell line were downloaded from GEO, accession number GSE221701, published by Agrawal-Singh et al [75]. H3K27me3 ChIP-seq peaks in .bed format from the HL-60 cell line were downloaded from GEO, accession number GSE175082, published by the ENCODE Project Consortium [76]. H3K27ac ChIP-seq peaks in .bed format from the HL-60 cell line were downloaded from GEO, accession number GSE167788, published by the ENCODE Project Consortium [76].

K562 Micro-C data in .mcool format was downloaded from GEO, accession number GSE206131, published by Barshad et al [77]. OCI-AML2 Hi-C data were downloaded in .fastq format from The European Genome-phenome Archive at the European Bioinformatics Institute, study ID EGAD00001006447, dataset ID EGAS00001004743, published by Takayama et al [31].

## Acknowledgements

This research was funded by Science Foundation Ireland through the SFI Centre for Research Training in Genomics Data Science under Grant number 18/CRT/6214 and supported in part by the EU’s Horizon 2020 research and innovation programme under the Marie Skłodowska-Curie grant H2020-MSCA-COFUND-2019-945385. Work in the Bond laboratory is supported by Science Foundation Ireland grants 20/FFP-P/8844 and 18/SPP/3522, the latter together with Children’s Health Ireland. Work in the Ryan laboratory is supported by Science Foundation Ireland grant 20/FFP-P/8641. Hi-C experiments were supported by a Haematology Association of Ireland Career Development Award to LJ. RL was supported by a DPhil in Cancer Sciences studentship awarded to by Cancer Research UK (Oxford Cancer Centre grant). AR is supported by a Wellcome Trust Clinical Research Career Development Fellowship (216632/Z/19/Z) and Medical Research Council (MRC, UK) Molecular Haematology Unit grant MC_UU_00029/7. The human fetal material was provided by the Joint MRC/Wellcome Trust Grant 099175/Z/12/Z Human Developmental Biology Resource (http://hdbr.org).

We thank Dr. Eric Conway, Dr. Aleksandar Krstic and members of the Bond, Ryan and Vaquerizas labs for useful suggestions on experiments and analyses.

## Data availability

Code to run RNA-Seq, CUT&RUN, ATAC-Seq, and part of Hi-C analyses is available at: https://github.com/cosmintudose/PRC2_AML_chromatin

Code to run Hi-C analyses was provided by the Vaquerizas lab and is partly based on: https://github.com/vaquerizaslab/Ing-Simmons_et_al_dorsoventral_3D_genome

