## Supplementary Figures S1-S5 for "Diverse epigenomic mechanisms underpin transcriptional dysregulation in Polycomb-altered acute myeloid leukemia"

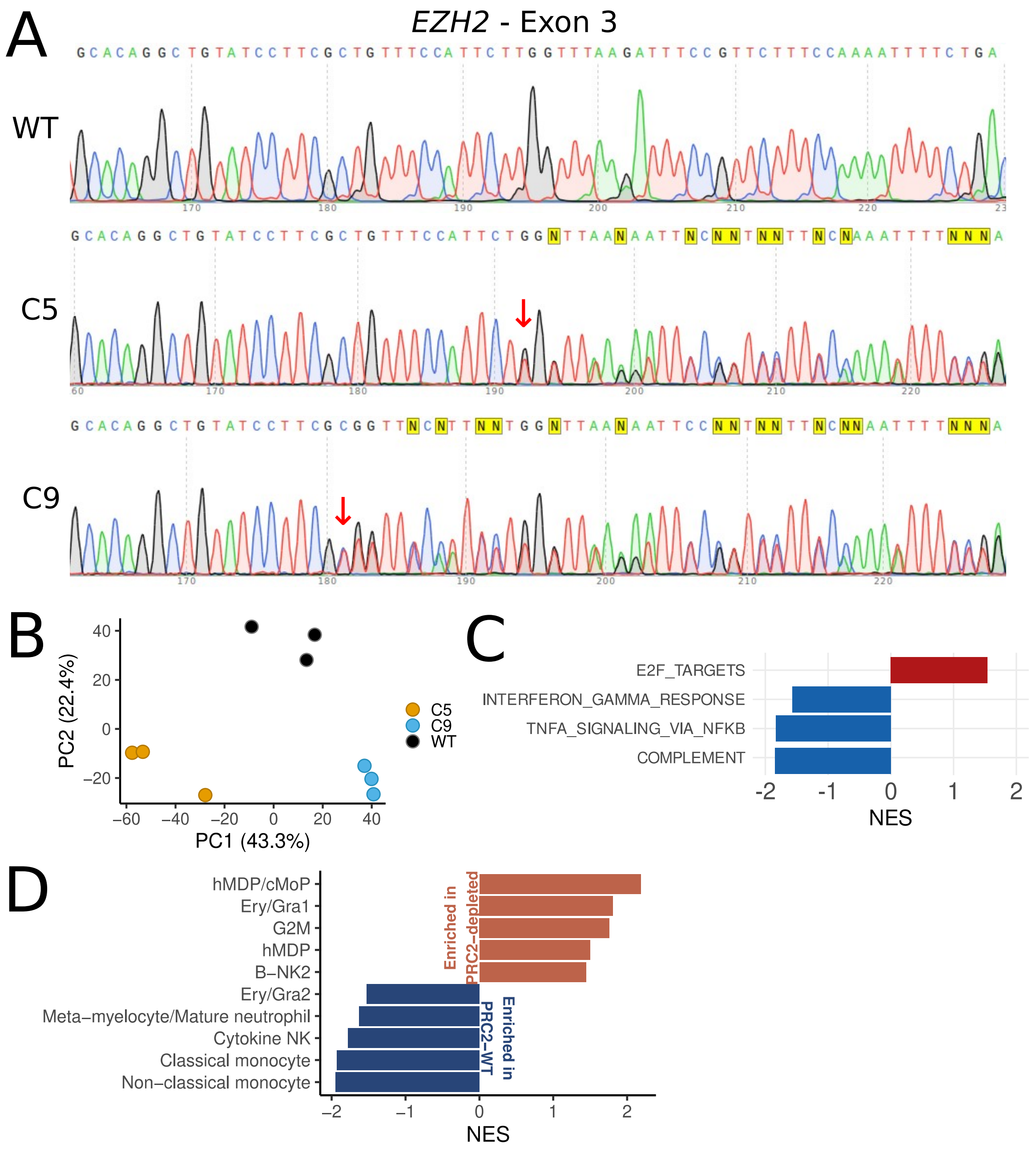


**Supplementary Figure S1 | Correlations between EZH2 depletion and genes involved in monocytic differentiation. A,** Sanger sequencing chromatogram showing CRISPR-Cas9 targeted region of *EZH2 (*exon 3) in WT, C5 and C9. **B,** PCA of OCI-AML2 RNA-Seq - all expressed genes. **C,** Gene set enrichment of MSigDB hallmarks comparing EZH2+/- (C5 and C9) and EZH2+/+ (WT) RNA-Seq. Red = Sets upregulated in EZH2+/- cells; blue = sets downregulated in EZH2+/- cells. **D,** GSEA performed on ranked differentially expressed genes in OCI-AML2 EZH2+/- cells vs EZH2+/+ cells using gene sets from the Human Blood Atlas, FDR = 5%.

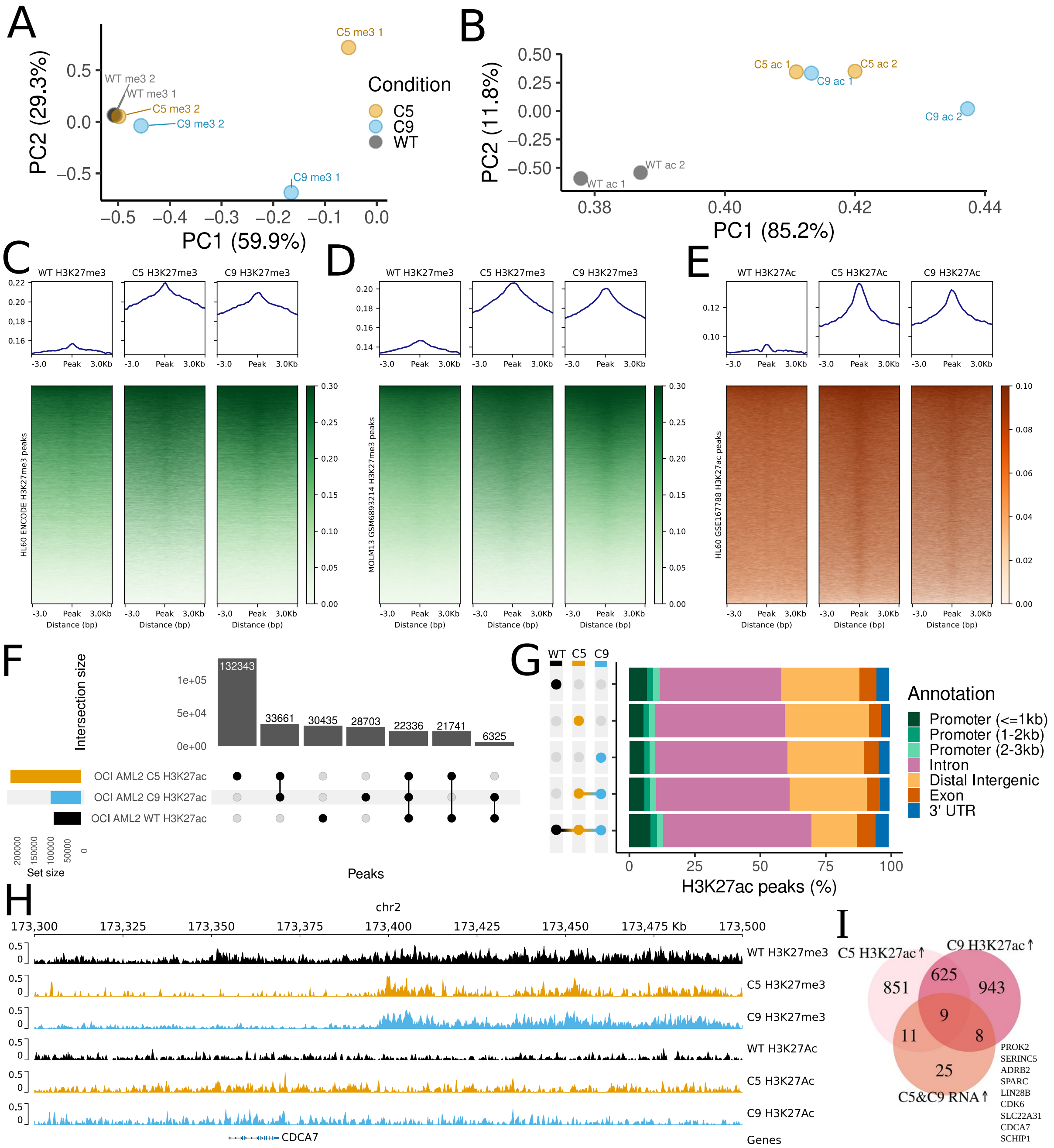


**Supplementary Figure S2 | Differences in H3K27me3 and H3K27ac between WT and EZH2-deficient clone C9. A, B,** PCA on binned bigwigs with bin size = 1kb of **(A)** H3K27me3 data and **(B)** H3K27ac data. **C, D,** WT C5 and C9 H3K27me3 signal at H3K27me3 peaks called in HL60 (**C**) and MOLM13 (**D**) from publicly available data (see Methods). **E,** WT C5 and C9 H3K27ac signals at H3K27ac peaks called in HL60 from publicly available data (see Methods). **F**, UpSet plot of peak counts for H3K27ac data. **G,** Annotation of H3K27ac peaks called in WT, C5 and C9. **H,** H3K27me and H3K27ac tracks for WT (black), C5 (orange) and C9 (cyan) at the *CDCA7* locus. **I,** Overlap between C5&C9 upregulated genes and genes that gain H3K27ac in C5 and C9; listed on the right the symbols of the genes overlapping in the three sets.

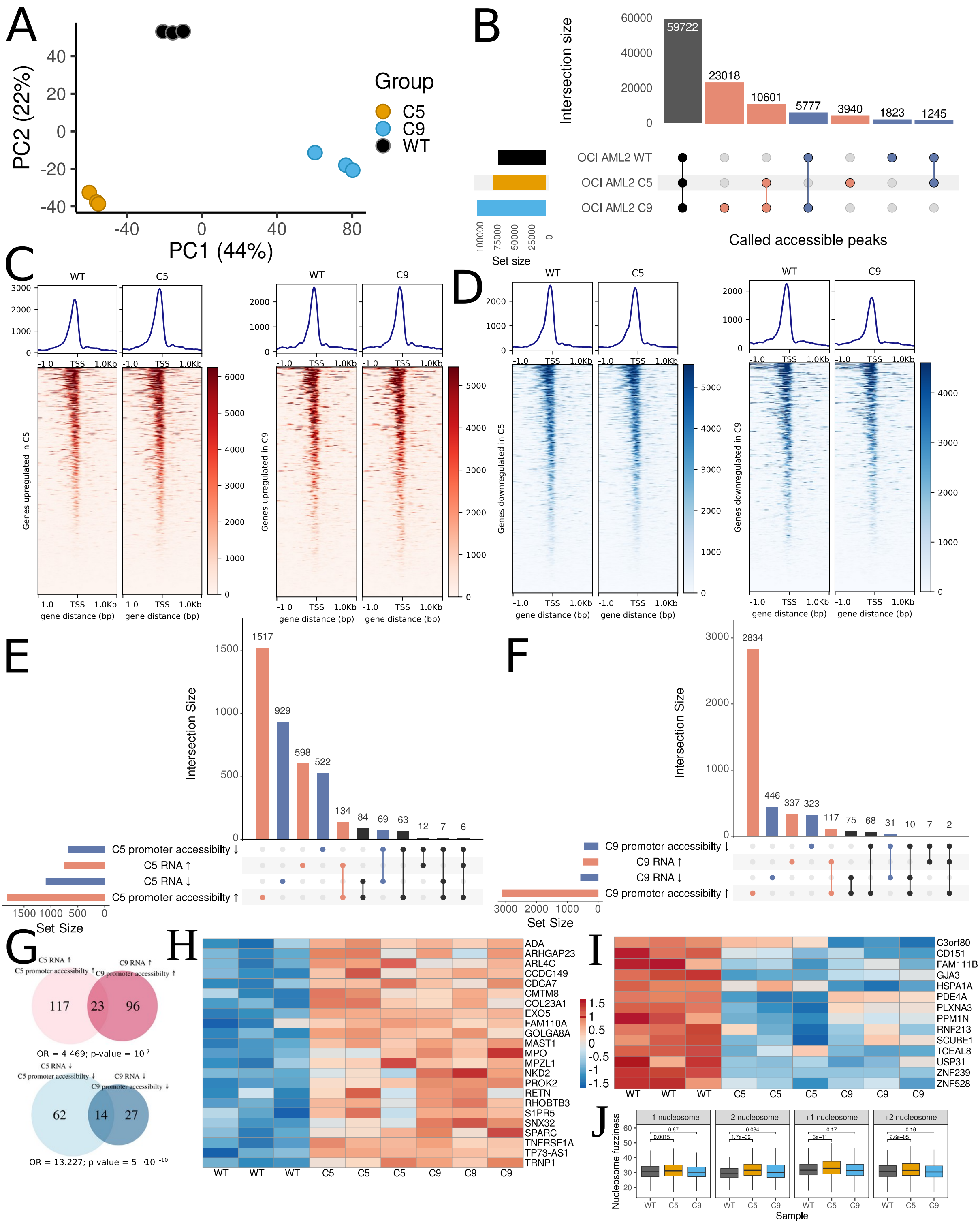


**Supplementary Figure S3 | ATAC-seq reveals changes in chromatin accessibility in WT, C5, and C9. A,** PCA of ATAC peaks from MACS2. **B,** Peak overlaps of peaks called by HMMRATAC. **C, D,** Nucleosome free region (NFR) signal at TSS of genes upregulated (**C**) and downregulated (**D**) in C5 and C9, respectively, compared with WT. **E, F,** Overlaps between genes with changed chromatin accessibility at the promoter and changed gene expression in C5 (**E**) and C9 (**F**) compared with WT. **G,** Overlaps of genes with increased promoter accessibility and increased RNA expression in C5 vs WT and C9 vs WT (red) and overlaps of genes with decreased promoter accessibility and decreased RNA expression in C5 vs WT and C9 vs WT. **H,** Genes with increased RNA expression and increased promoter accessibility in both C5 and C9 vs WT. **I,** Genes with decreased RNA expression and decreased promoter accessibility in both C5 and C9 vs WT. **J,** Nucleosome fuzziness at -2, -1, +1 and +2 nucleosomes for WT, C5 and C9. Box = 1st and 3rd quartiles; middle black line = median; whiskers extend to 95% of data points. Values above brackets indicate p-values from Wilcoxon unpaired tests.

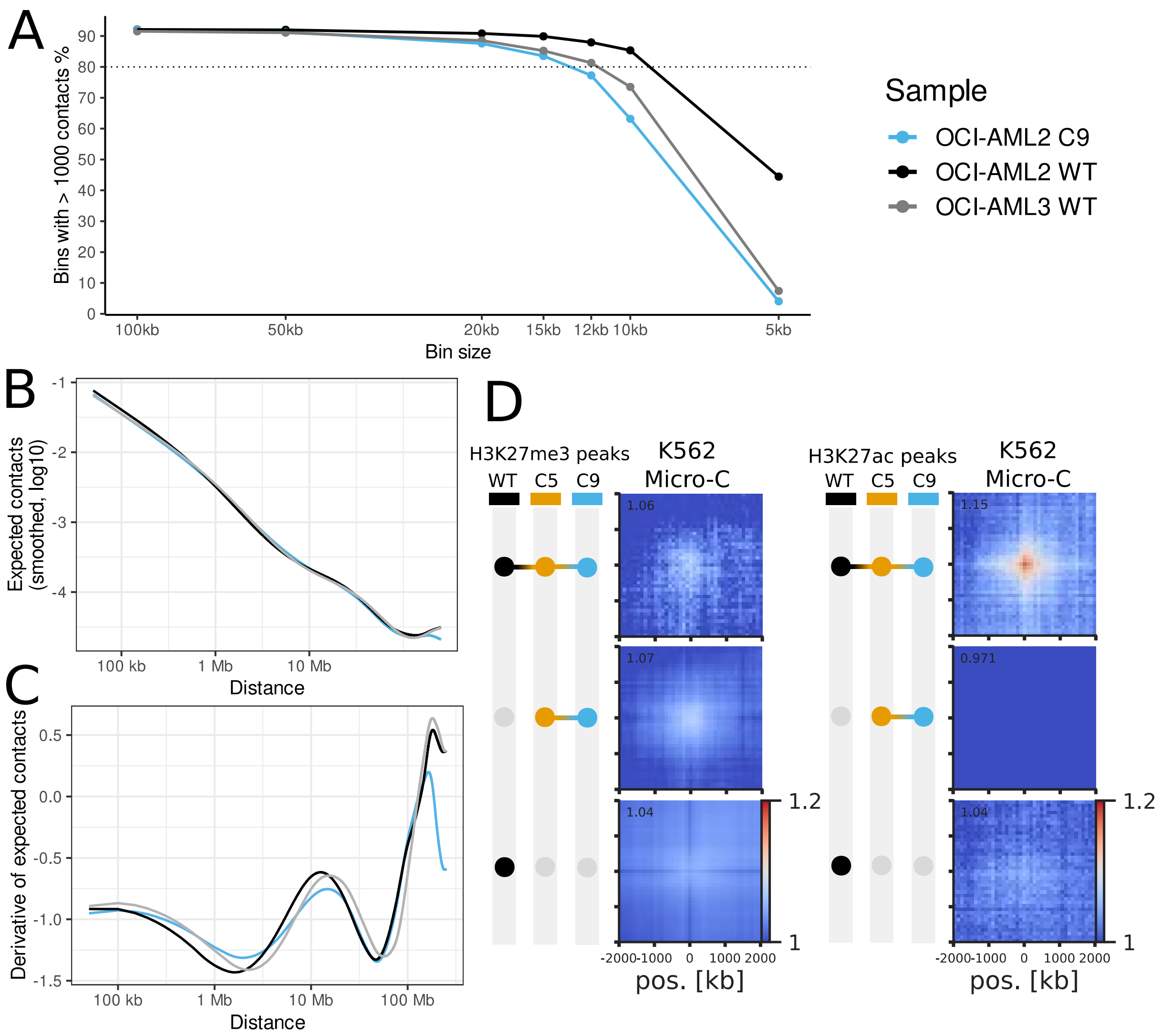


**Supplementary Figure S4 | QC on Hi-C data and validation on Micro-C. A,** Resolution of Hi-C data for each sample (see Methods). **B,** Distance-decay curve for each sample **C,** Derivative of the distance-decay curve for each sample **D,** Pileups of contacts between CUT&RUN H3K27me3 and H3K27ac peaks in Micro-C data from K562 at 100kb resolution and 2Mb flanking regions.

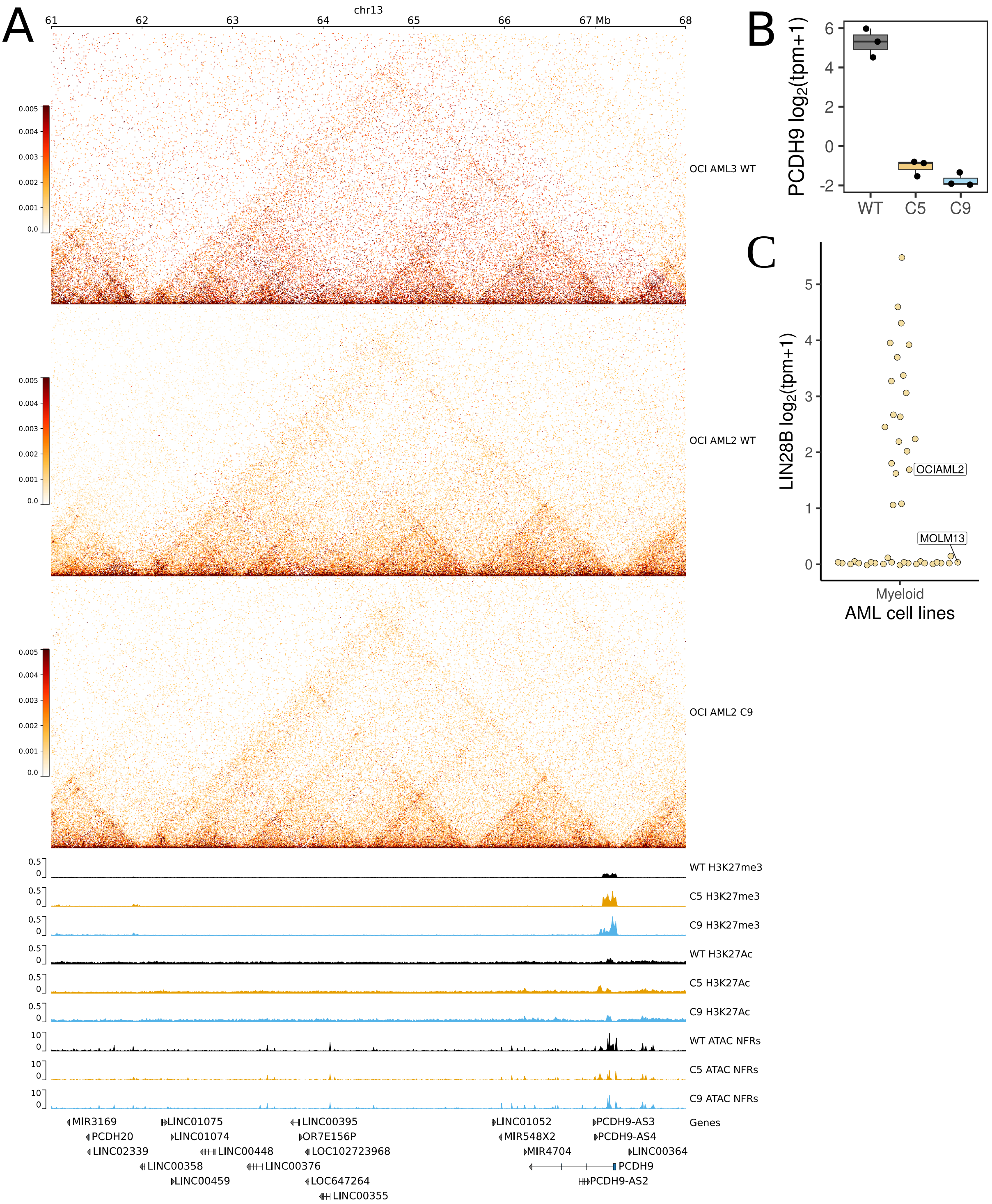


**Supplementary Figure S5 | Integration of epigenomic analysis results at selected loci.** **A,** Hi-C (OCI-AML3, OCI-AML2 WT and C9), H3K27me3 and H3K27ac and ATAC NFRs (WT - black, C5 - orange and C9 - cyan) tracks at PCDH9 **B,** Expression of *PCDH9* in WT, C5 and C9. **C,** LIN28B expression in the Cancer Cell Line Encyclopaedia (CCLE) in AML cell lines (N = 43).

**Supplementary Table S1 | List of antibodies**

| Protein | Supplier | Catalogue Number | Species | Dilution |
|---|---|---|---|---|
| EZH2 (D2C9) XP® Rabbit mAb #5246 | Cell Signaling Technology | #5246 | Rabbit | 1:1000 |
| Tri-Methyl-Histone H3 (Lys27) (C36B11) Rabbit mAb #9733 | Cell Signaling Technology | #9733 | Rabbit | 1:1000<br>1:100 |
| Acetyl-Lysine H3 (Lys27) (D5E4) XP Rabbit mAb #8173 | Cell Signaling Technology | #8173 | Rabbit | 1:1000<br>1:100 |
| GAPDH (14C10) Rabbit mAb #2118L | Cell Signaling Technology | #2118L | Rabbit | 1:1000 |
| IgG XP® Isotype Control (DA1E) XP® Rabbit mAb | Cell Signaling Technology | #3900S | Rabbit | 1:100 |

**Supplementary Table S2 | Differential expression analysis OCI-AML2 EZH2+/- vs EZH2+/+**

**Supplementary Table S3 | GSEA on cell lines using ABC gene sets**

**Supplementary Table S4 | Annotated H3K27me3 and H3K27ac called peaks in WT, C5 and C9**

**Supplementary Table S5 | Annotated ATAC open chromatin regions in WT, C5 and C9**

**Supplementary Table S6 | GO:BP enrichment with GREAT tool on regions more accessible in clones, compared to WT**
